# RhoA Allosterically Activates Phospholipase Cε via its EF Hands

**DOI:** 10.1101/2024.11.14.623250

**Authors:** Vaani Ohri, Kadidia Samassekou, Kaushik Muralidharan, Elisabeth E. Garland-Kuntz, Isaac J. Fisher, William C. Hogan, Bailey M. Davis, Angeline M. Lyon

**Author notes:** To whom correspondence should be addressed: Angeline M. Lyon, Departments of Chemistry and Biological Sciences, Purdue University, 560 Oval Drive, West Lafayette, Indiana 47907. Karp Building, Room RB08004, 1 Blackfan Circle, Boston, MA 02115. Sterling Hall of Medicine, Yale University, 333 Cedar St, New Haven, CT 06510. Abigail Wexner Research Institute at Nationwide Children’s, Research Building II, WA2106, Columbus, OH 43205.

## Abstract

Phospholipase Cε (PLCε) cleaves phosphatidylinositol lipids to increase intracellular Ca^2+^ and activate protein kinase C (PKC) in response to stimulation of cell surface receptors. PLCε is activated via direct binding of small GTPases at the cytoplasmic leaflets of cellular membranes. In the cardiovascular system, the RhoA GTPase regulates PLCε to initiate a pathway that protects against ischemia/reperfusion injuries, but the underlying molecular mechanism is not known. We present here the cryo-electron microscopy (cryo-EM) reconstruction of RhoA bound to PLCε, showing that the G protein binds a unique insertion within the PLCε EF hands. Deletion of or mutations to this PLCε insertion decrease RhoA-dependent activation without impacting regulation by other G proteins. Together, our data support a model wherein RhoA binding to PLCε allosterically activates the lipase and increases its interactions with the membrane, resulting in maximum activity and cardiomyocyte survival.

## INTRODUCTION

Mammalian phospholipase C (PLC) enzymes are translocated and activated at the cytoplasmic leaflet of membranes in response to diverse stimuli. All PLCs cleave phosphatidylinositol-4,5-bisphosphate (PIP_2_) at the plasma membrane, producing inositol-1,4,5-triphosphate (IP_3_) and diacylglycerol (DAG). IP_3_ stimulates intracellular Ca^2+^ release, which together with DAG, activates protein kinase Cs (PKCs)^1^. PLCε also cleaves phosphatidylinositol-4-phosphate (PI4P) at the perinuclear membrane, where local increases in DAG activate PKCs and protein kinase D^2,3^.

PLCε is activated downstream of G protein-coupled receptors (GPCRs) and receptor tyrosine kinases (RTKs) through binding of the Rap1A, RhoA, and Ras GTPases, as well as the Gβγ heterodimer. Activation likely proceeds through simultaneous membrane localization and activation, with the G protein dictating the location of activation^2–4^. In the cardiovascular system, PLCε activation has been best studied in response to stimulation of G_s_- and G_12/13_-coupled receptors. Stimulation of the β-adrenergic receptors (β-AR) leads to activation of the Rap1A GTPase, which in turn activates PLCε at the perinuclear membrane. Increased PI4P hydrolysis activates PKC- and PKD-dependent pathways that maximize Ca^2+^-induced Ca^2+^ release (CICR) and contractility^5,6^. However, sustained activation leads to upregulation of genes that promote cardiac hypertrophy^7–11^. Gβγ-dependent activation of PLCε, downstream of the endothelin-1 receptor, results in a similar pathological response^10–12^. Intriguingly, PLCε has a cardioprotective role in response to ischemia/reperfusion injuries. Stimulation of G_12/13_-coupled receptors, including the sphingosine-1-phosphate receptor (S1PR), activates RhoA^2,3^. In this pathway, RhoA activates PLCε at the plasma membrane, where PIP_2_ hydrolysis increases intracellular Ca^2+^ and PKC activity. The latter activates a PKD-dependent pathway that protects the mitochondria from oxidative stress, a major cause of acute cardiomyocyte cell death under ischemic conditions^13–17^.

The ability of PLCε to hydrolyze substrates at different intracellular sites is due to its subfamily-specific regulatory domains and insertions. Like other PLCs, PLCε contains a pleckstrin homology (PH) domain, four EF hand repeats (EF1-4), the catalytic TIM barrel, and C2 domain (**Fig. 1a**)^3,18^. The core is flanked by an N-terminal region and a CDC25 domain that is a guanine nucleotide exchange factor (GEF) for the Rap1A GTPase. At its C-terminus are two Ras association (RA) domains, RA1 and RA2. RA1 stabilizes the catalytic core, while RA2 binds Rap1A and Ras GTPases^3,19,20^. Finally, the TIM barrel contains two insertions: the X–Y linker and the Y-box. As in PLCβ and PLCδ, the PLCε X–Y linker occludes the active site and must be displaced via an interfacial activation mechanism to allow substrate binding^3,21,22^. The Y-box is a ∼70 amino acid insertion unique to the PLCε subfamily^23^, but whether it has a role in basal activity less clear.

**Figure 1.**
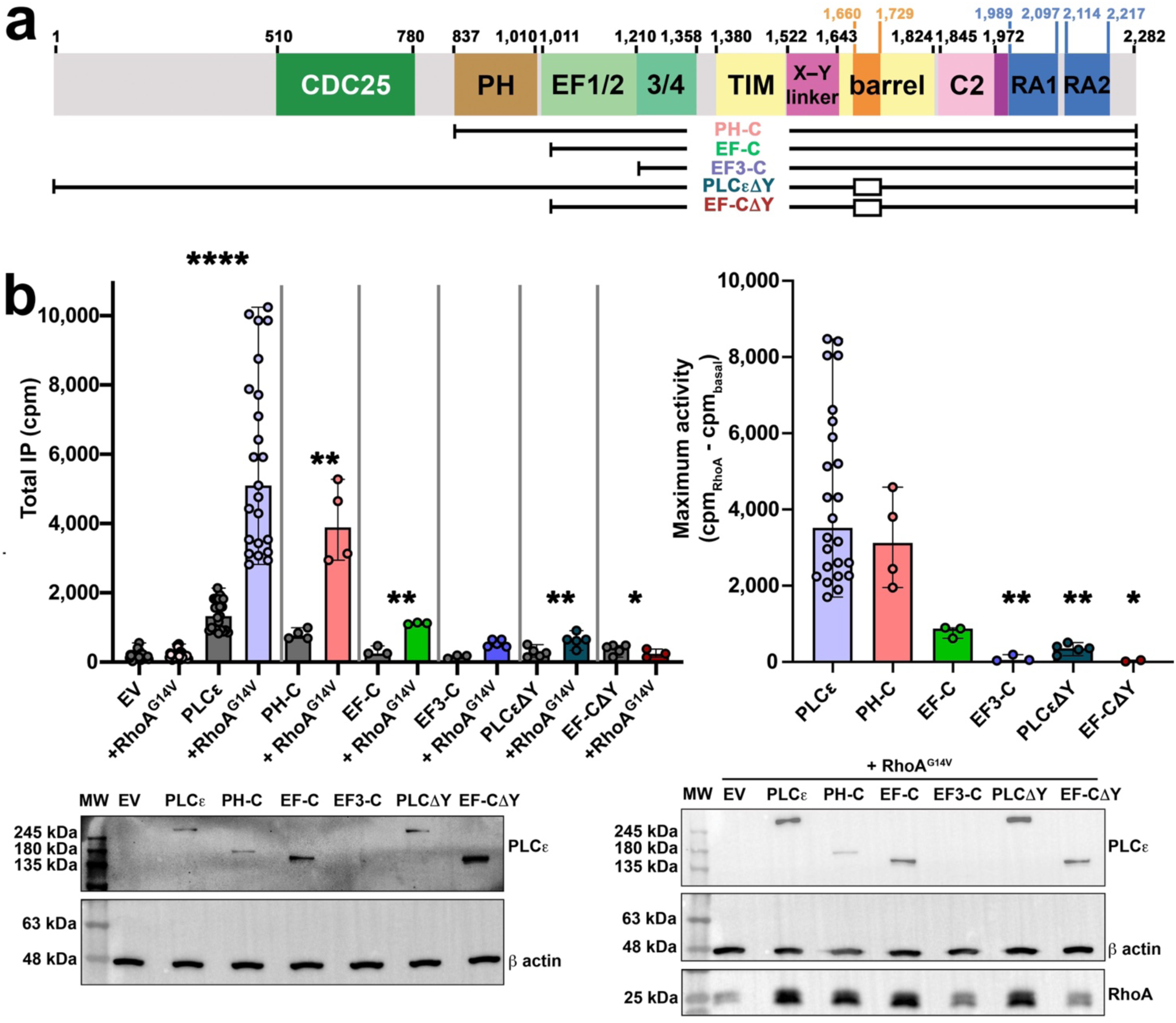
The PLCε EF1/2 hands and Y-box are required for maximum RhoA-dependent activity. **(a)** Domain architecture of *R. norvegicus* PLCε, with domain boundaries shown above. Variants used in this study are shown below, open boxes indicate internal deletions. **(b)** (*Left*) Basal and RhoA^G14V-^ stimulated activities of PLCε and variants retaining EF1/2 and/or the Y-box. At least three independent experiments from independent transfections were carried out for each variant, and data is shown as the average of triplicate measurements ± SD. Data was analyzed using an unpaired, one-tailed t-test with Welch’s correction comparing the basal and RhoA-stimulated activities of each variant. ****p<0.0001, for PH-C **p<0.0053, EF-C **p<0.0034, PLCε1Y **p<0.0075, EF-C1Y *p<0.012. (*Right*) The change in maximal activity ± SD was calculated by subtracting RhoA^G14V^-stimulated activity by the basal activity of each variant. Data was analyzed using a one-way ANOVA and Kruskal-Wallis test comparing each variant to PLCε, followed by a Dunn’s multiple comparisons test. For EF3-C **p<0.0053, PLCε1Y **p<0.0030, and EFC-1Y *p>0.0159. Representative western blots are shown below, with empty pCMV vector (EV) and β-actin used as loading controls. Differences in expression were not found to be statistically significant, but may still contribute to variation in activities. PLCε variants express a C-terminal FLAG tag and are detected with an anti-FLAG antibody, while RhoA contains an N-terminal HA tag and is detected with an anti-HA antibody.

RhoA is the most robust activator of PLCε, increasing its activity ∼5-10-fold over basal in cell-based assays^3,24,25^. Initial studies demonstrated that only the active form the GTPase directly interacted with the lipase to increase activity. Efforts to map its binding site relied on a series of N- and C-terminally truncated PLCε variants that narrowed its binding site to a region between the EF hands and C2 domains (**Fig. 1a**). Given that only PLCε contains the Y-box, it emerged as a possible binding site for the GTPase, especially because its deletion eliminated RhoA-dependent activation. However, N-terminally truncated PLCε variants, with or without a Y-box, were shown to pull down the active GTPase to similar extents^23,26^. Although the Y-box may be required for RhoA-mediated activation, it is clearly not the binding site.

In this work, we define the minimal structural requirements for RhoA-dependent activation of PLCε. Prenylated RhoA×GTP is required for maximum activation, but a soluble mutant also stimulates the lipase, demonstrating the mechanism likely involves both membrane localization and allosteric components. Informed by newly annotated PLCε domain boundaries, we show PLCε variants retaining the PH domain and EF hands 1/2 (EF1/2) are robustly activated by the GTPase, whereas variants that lack the Y-box, the N-terminus, CDC25, PH domain, and/or EF1/2 hands decreased basal and RhoA-stimulated activities. Our cryo-electron microscopy (cryo-EM) reconstruction of a RhoA×GTP–PLCε complex reveals an integral role of the EF hands in the mechanism, because RhoA×GTP binds to a PLCε subfamily-specific insertion in this domain, ∼60 Å away from the active site and Y-box in the TIM barrel. Mutation or deletion of the PLCε EF hand insertion compromises RhoA-dependent activation in cells. Comparison of the RhoA×GTP–PLCε reconstruction to other PLCε structures shows that RhoA binding induces conformational changes within the EF hands that likely contribute to allosteric activation.

## RESULTS AND DISCUSSION

### RhoA-dependent activation of PLCε requires EF hands ½

Previous studies investigating RhoA-dependent activation of PLCε used variants truncated at the N-terminus, removing all or parts of the CDC25 domain, PH domain, and EF1/2, as based on sequence conservation, or the C-terminal RA domains^14,27,28^. Because we recently established domain boundaries for the PH and EF hand domains^18^, we used this approach to test the contributions of the N-terminal regions and Y-box in basal and RhoA-dependent activation in cells. In this assay, cells are metabolically labelled with [^3^H]-myoinositol which is incorporated into their lipid head groups. Transfected PLCε species cleave the [^3^H]-labelled phosphatidylinositol phospholipids, producing DAG and [^3^H]-inositol species ([^3^H]-IPx), the latter of which are quantified by scintillation counting^29,30^. RhoA^G14V^ increases WT PLCε activity ∼5-fold over basal, consistent with previous reports (**Fig. 1a,b**)^23,31^. PLCε PH-C, which lacks the N-terminal 836 residues, is similarly activated by RhoA^G14V^. PLCε EF-C, which lacks the N-terminus and PH domain, is also significantly activated by RhoA^G14V^, but its maximum basal activity is ∼4-fold lower than that of RhoA-activated PLCε. PLCε EF3-C, which is further truncated to remove EF1/2, is unresponsive to the GTPase (**Fig. 1a,b**). We also reassessed the role of the Y-box in activation in the context of PLCε and the EF-C variant. Deletion of this element significantly decreased basal and eliminated RhoA stimulation (**Fig. 1b**), indicating the Y-box is required for lipase activity in general, and not specifically for activation by RhoA.

Maximum RhoA-dependent activity of PLCε variants decreased progressively as its N-terminus was truncated. To confirm the role of the EF hands, particularly EF1/2, in RhoA activation of the holoenzyme, we generated a series of chimeras between PLCε and PLCβ3, which is not regulated by RhoA (**Fig. 2a**). In these chimeras, either the entire EF1-4 module of PLCε was replaced with that of PLCβ3 (PLCε/β3 EF), or only the EF1/2 (PLCε/β3 EF1/2) or EF3/4 (PLCε/β3 EF3/4) module was replaced (**Fig. 2a)**. The chimeras expressed similarly and were properly folded and functional (**Supplemental Fig. 1**). However, only PLCε/β3 EF1/2 showed significantly activation by RhoA^G14V^, with a ∼3-fold increase over basal activity (**Fig. 2b**).

**Figure 2.**
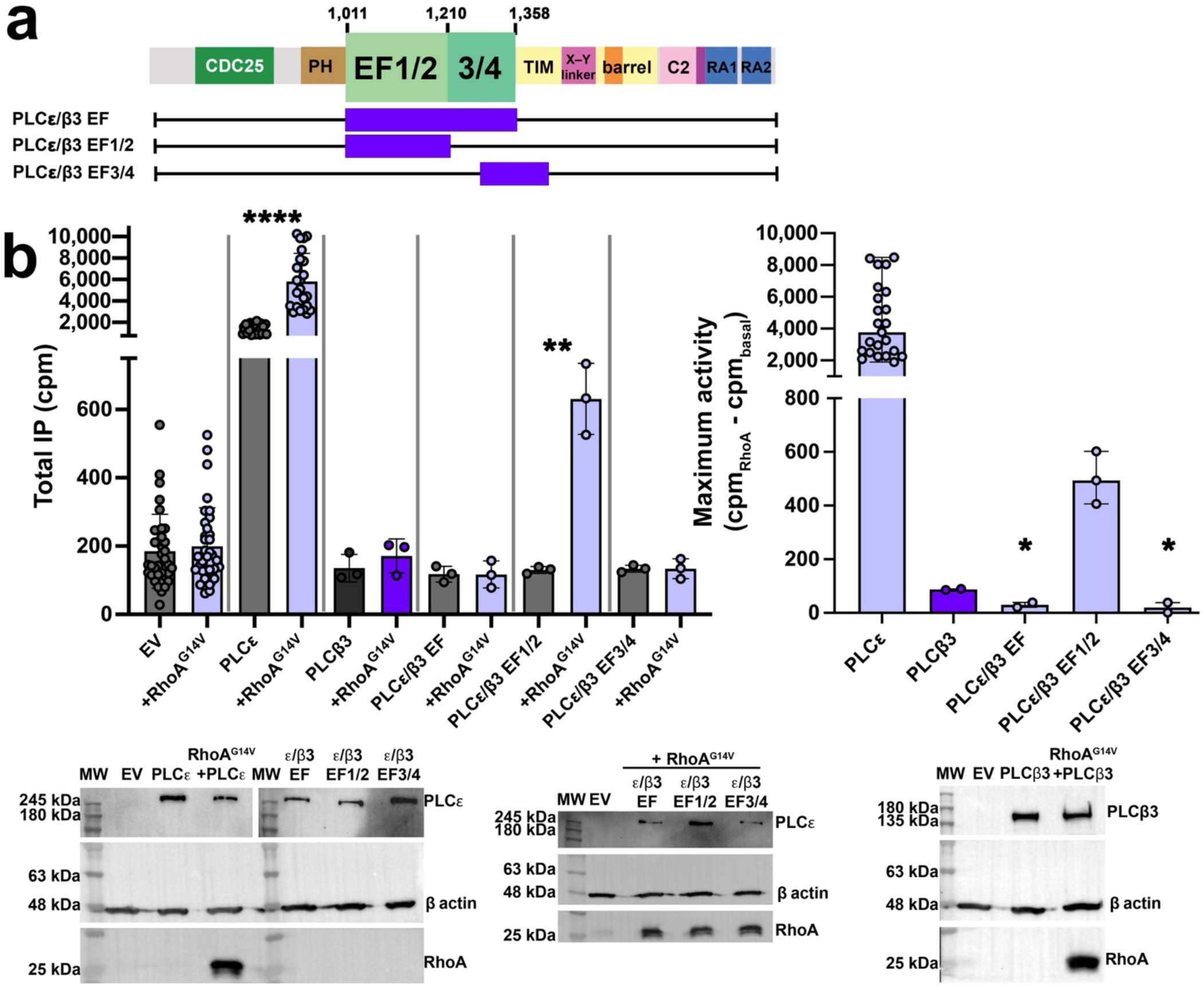
PLCε/β3 chimeras confirm the EF hands are essential for RhoA-dependent activation. **(a)** Schematic of the PLCε/β3 EF hand chimeras. **(b)** Basal and RhoA^G14V^-stimulated activities of PLCε, PLCβ3, and PLCε/β3 chimeras. (*Left*) Only PLCε variants that retain the EF1/2 hands are responsive to RhoA-dependent activation. At least three independent experiments from independent transfections were performed for each variant, and data is shown as the average of triplicate measurements ± SD. Data was analyzed using an unpaired, one-tailed t-test with Welch’s correction comparing the basal and RhoA-stimulated activities of each variant. ****p<0.0001, **p<0.0067, *p<0.0167. (*Right*) Changes in maximal activity of each variant were analyzed using a one-way ANOVA and Kruskal-Wallis test comparing each variant to PLCε, followed by a Dunn’s multiple comparisons test. ***p<0.0485, *p<0.0198. Representative western blots are shown below, with empty pCMV vector (EV) and β-actin used as loading controls. PLCε variants express a C-terminal FLAG tag and are detected with an anti-FLAG antibody, and RhoA contains and N-terminal HA tag for detection using an anti-HA antibody.

### Maximum activity requires prenylated RhoA

For PLC enzymes regulated by G proteins, maximum lipase activation requires the G protein activators to be prenylated and/or acylated^3,32^. We compared the ability of wild-type, prenylated RhoA×GTPγS and soluble forms of RhoA×GTPγS and RhoA^G14V^×GTP to directly activate purified PLCε PH-C using a modified version of the commercially available IP-One assay. Briefly, phosphatidylinositol (PI) is incorporated into liposomes, and the activity of the lipase produces DAG, which remains in the liposome, and free inositol phosphate (IP1)^33,34^. Wild-type (prenylated) RhoA×GTPγS and soluble RhoA^G14V^×GTP significantly increased lipase activity ∼7 and ∼5-fold over basal, respectively. Soluble RhoA×GTPγS increased lipase activity ∼3-fold over basal (**Supplemental Fig. 2**). The fact that soluble RhoA variants partially activate PLCε PH-C confirms that membrane localization mediated by RhoA alone is insufficient to achieve full activation of the lipase by RhoA.

### RhoA×GTP binds to the PLCε E2α’ helix of the EF hands

Cryo-EM single particle analysis (SPA) was used to determine the structure of RhoA×GTP bound to PLCε PH-C. This variant is activated by RhoA (**Fig. 1a,b**) and is the largest variant that has been purified in sufficient quantities for biophysical studies^18,30,35^. PLCε PH-C and prenylated RhoA×GTP were incubated in a 3:1 molar ratio before being applied to grids. From an initial data set of 1,329,298 particles, two distinct populations containing 184,875 and 106,370 particles were identified. Because the first population was larger and the resulting volume showed more structural features, it was selected for further processing and used to generate an initial 3.5 Å map. A structure of PLCε PH-C (PDB ID 9B13, *in review*^18^) was fit into the map, revealing unmodeled density around the EF hands. Given the importance of the EF hands for RhoA-dependent activation (**Fig. 1, 2**), the crystal structure of RhoA×GMPPNP (PDB ID 1S1C)^36^ was placed in the density such that its switch regions, which had the strongest density, were adjacent to the EF hands. The resulting RhoA–PLCε PH-C model was then used as template for additional rounds of particle picking and refinement, resulting in a final map at 3.3 Å resolution (209,463 particles, **Supplemental Figs. 3-5, Supplemental Table 1**).

In the RhoA×GTP–PLCε PH-C reconstruction, clear density is observed for the PH-RA1 domains (**Fig. 3**). No density is observed for the RA2 domain, despite its presence in the protein because it is flexibly connected to the rest of the lipase^19,30^. The architecture of PLCε is similar to that of the lipase bound to an Fab fragment, with an r.m.s.d. of 0.87 Å for 786 Cα atoms (out of 841 resolved residues, PDB ID 9B13)^18^. The PH domain and EF1/2 pack adjacent to the PLCε catalytic core, which includes the EF3/4-RA1 domains^35^. The active site remains blocked by the C-terminus of the X–Y linker. However, the rest of the X–Y linker (residues 1525-1631) and the Y-box (residues 1662-1730) were not resolved, consistent with these regions being highly dynamic.

**Figure 3.**
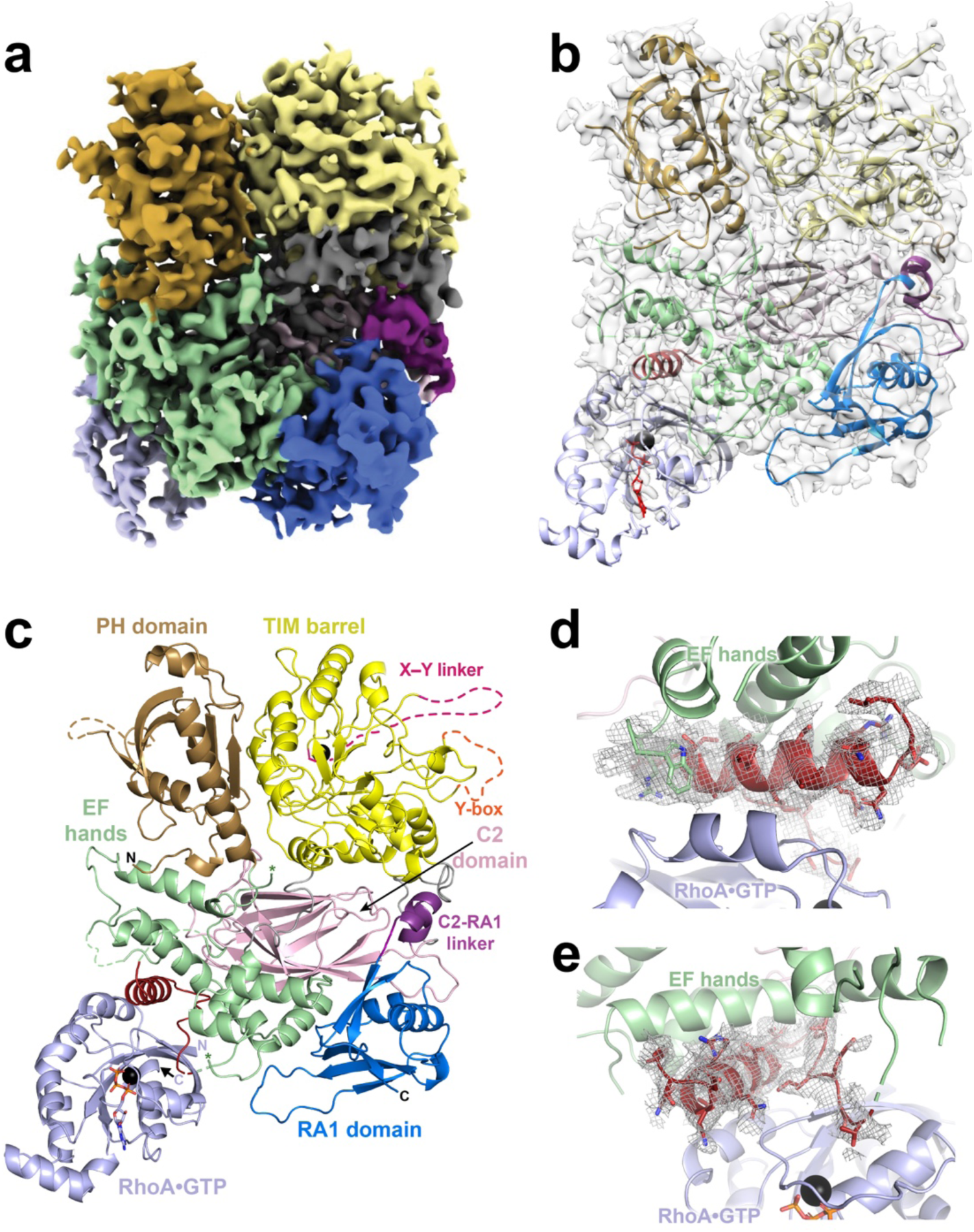
RhoA·GTP binds to the PLCε EF hands. (**a**) The 3.3 Å cryo-EM map of the RhoA·GTP–PLCε PH-C complex and (**b**) fit with the ribbon diagram of the RhoA·GTP–PLCε PH-C complex. The domains in PLCε are colored as in Fig. 1A and RhoA·GTP is shown in light blue. (**c**) Ribbon diagram of the RhoA·GTP–PLCε PH-C colored as in (a). The PLCε active site Ca^2+^ is shown as a black sphere, the E2α’ helix and adjacent loop that bind RhoA are shown in firebrick. Disordered regions, including the X–Y linker (hot pink) and Y-box (orange) are shown as dashed lines. RhoA·GTP is shown in light blue, Mg^2+^ as a black sphere, and GTP in light blue sticks. The N- and C-termini of each protein are labelled. (**d**) Density (gray mesh) for the E2α’ helix (shown in red) and (**e**) loop connecting the helix to the EF3/4 subdomain.

RhoA×GTP binds exclusively to the EF hands via its switch I and switch II, consistent with other RhoA–effector enzyme complexes (**Fig. 3, Supplemental Fig. 6)**^36–38^. Switch I and II of RhoA×GTP could be resolved in the density (Supplemental Fig. 7) and bind to the PLCε E2α’ helix (residues 1273-1287) in a subfamily-specific insertion within EF3/4^18,39^. The E2α’ helix is followed by an extended loop (residues 1288-1302) that reenters the EF3/4 module. Residues 1288-1296 are ordered and poised to interact with residues on the β3 strand of the GTPase (**Figs. 3d, 4; Supplemental Fig. 7**). Overall, the interaction is largely hydrophobic, burying ∼1,800 Å^2^ surface area. Within the PLCε E2α’ helix, residues Asn1275, Ile1279, Ala1282, Ile1283, and Ala1286 interact with Phe39 in switch I and Leu69 and Leu72 in switch II of RhoA×GTP (**Figs. 3d, 4)**. PLCε Ile1295, on the loop following E2α’, also interacts with RhoA Phe39. Finally, PLCε Arg1049 and Trp1051, located on the loop connecting EF1 and EF2, interact with Leu72 and Pro75 at the C-terminus of the RhoA×GTP switch II helix (**Figs. 3d, 4**). The orientation of RhoA×GTP and PLCε places the prenylated C-tail of the GTPase in the same plane as the PLCε PH domain and the active site in the TIM barrel, which would allow these elements to simultaneously engage the membrane as expected during activation^18,35^.

**Figure 4.**
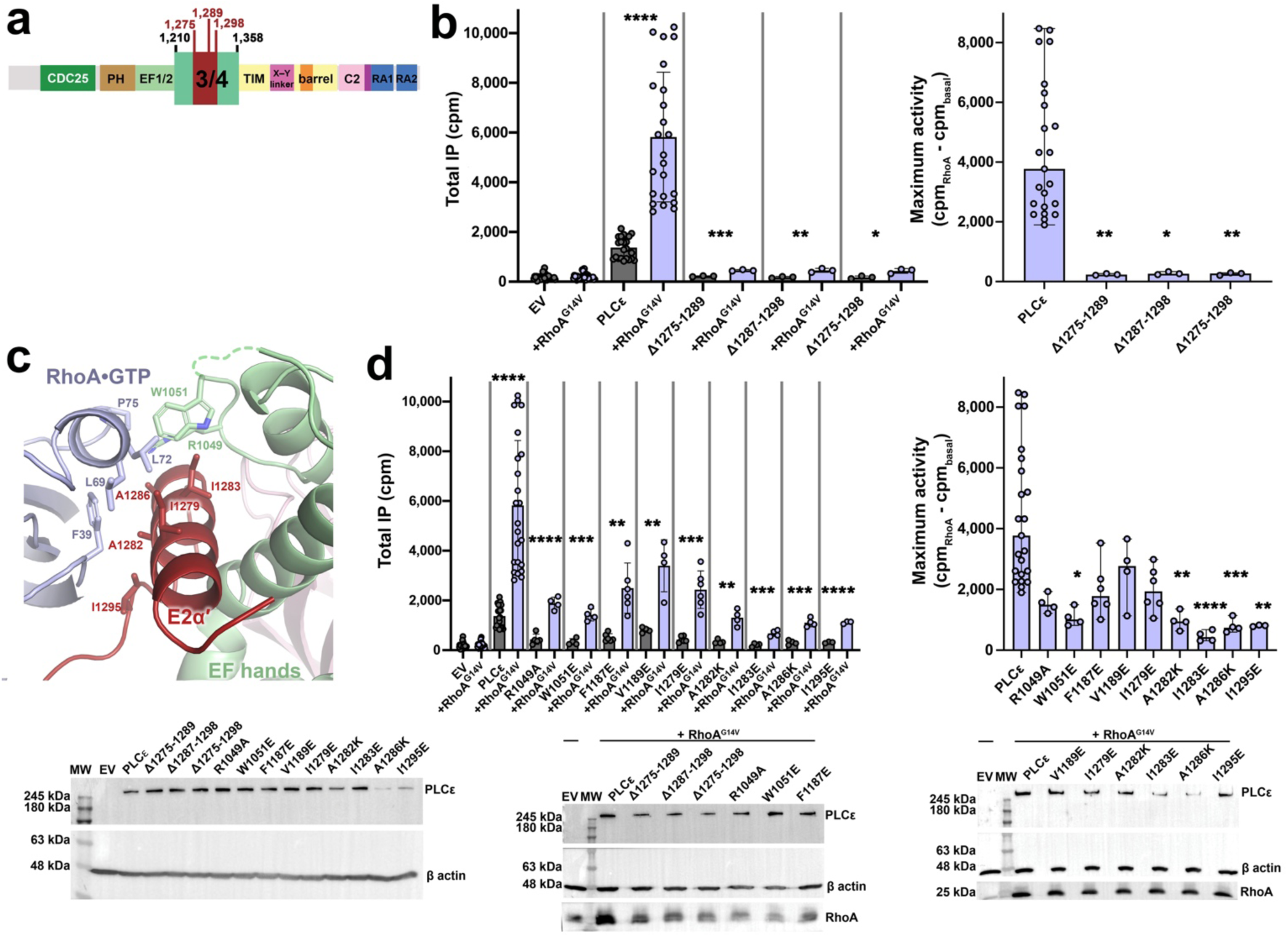
The PLCε E2α’ helix is needed for RhoA-dependent activation. **(a)** Schematic showing the boundaries for internal deletions in the PLCε EF3/4 module. (**b**) (*Left*) Basal and RhoA^G14V^-stimulated activities of PLCε variants lacking the E2α’ helix (Δ1275-1289), the loop connecting it to the F3α helix (Δ1287-1298) in the EF3/4, or both (Δ1275-1298). Deletion of any of these regions largely eliminates RhoA-dependent activation. At least three independent experiments from independent transfections were performed for each variant. Data shown represents the average of triplicate measurements ± SD, and analyzed using unpaired, one-tailed t-test with Welch’s correction to compare the basal and RhoA-stimulated activities of each variant. ****p<0.0001, ***p<0.0003, **p<0.0080, *p<0.0112. (*Left*) The change in maximal activity ± SD was calculated by subtracting RhoA-stimulated activity by the basal activity of each variant. Data was analyzed using a one-way ANOVA and Kruskal-Wallis test comparing each variant to PLCε, followed by a Dunn’s multiple comparisons test. For Δ1275-1289, **p<0.0041, for Δ1287-1298, *p<0.0134, and for Δ1275-1298, **p<0.0097. **(c)** The PLCε E2α’ helix (red) binds to the switch regions of RhoA (light blue). Additional contacts with RhoA are made by residues in the EF1/2 module and the loop linking E2α’ to the F3α helix. Labelled residues were subjected to site-directed mutagenesis and their impact on RhoA-dependent activation quantified. **(d)** (*Left*) Mutations in the G protein–PLCε interface decrease RhoA-dependent activation. At least three independent experiments from independent transfections were carried out for each variant. Data shown represents the average of triplicate measurements ± SD, and was analyzed using unpaired, one-tailed t-test with Welch’s correction to compare the basal and RhoA-stimulated activities of each variant. ****p<0.0001, ***p<0.0003, **p<0.0080, *p<0.0112. (*Right*) Mutation of PLCε Trp1051 in EF1/2, residues Ala1282, Ile1283, Ala1286 in E2α’, and Ile1295 in the E2α’-F3α loop significantly decrease maximum RhoA-dependent activation. Data was analyzed using a one-way ANOVA and Kruskal-Wallis test comparing each variant to PLCε, followed by a Dunn’s multiple comparisons test. Representative western blots are shown below, with empty pCMV vector (EV) and β-actin used as loading controls. Differences in expression were not found to be statistically significant, but may still contribute to variation in activities PLCε variants express a C-terminal FLAG tag and are detected with an anti-FLAG antibody, while RhoA contains an N-terminal HA tag, and detected using an anti-HA antibody.

There are differences in the intramolecular interactions between the RhoA complex and the PLCε EF3-RA1 and Fab–PLCε PH-C structures^18,30^. Residue-residue (RR) distance analysis^40^ was used to identify conformational changes across the three structures in an unbiased manner (**Supplemental Figs. 8, 9**). Within the individual structures, the PH and TIM barrel domains move as a single unit, as do EF3/4 and the C2 domain. In contrast, the RA1 domain is conformationally distinct in each structure, potentially due to the influence of crystal packing interactions in the EF3-RA1 structure or Fab binding in the Fab– PLCε PH-C structure (**Supplemental Figs. 8, 9**)^18,30^. Binding of RhoA×GTP to the EF hands induces a shift in the position of EF3/4, moving it ∼2 Å closer to the TIM barrel domain relative to its position in the other structures. This conformational change may reflect an allosteric component of RhoA-mediated activation, given its potential impact on the catalytic TIM barrel domain (**Supplemental Fig. 10).**

### The PLCε E2α’ helix is required for RhoA×GTP-dependent activation

We first tested whether the PLCε E2α’ helix and flanking loop are needed for RhoA-dependent activation (**Fig. 4a,b, Supplemental Figure 7)**. Deletion of the disordered region preceding the E2α’ helix (PLCε Δ1226-1270) did not eliminate RhoA-dependent activation but decreased its maximum activity (**Fig. 4a,b**). Deletions of the E2α’ helix (Δ11275-1289), the flanking loop (Δ1287-1298), or both (Δ1275-1298) eliminated RhoA-dependent activation. The E2α’ helix is only required for activation by RhoA, as PLCε and PLCε Δ11275-1289 had the same fold activation when co-transfected with two other well-established G protein activators, Rap1A^Q63E^ and the Gβγ heterodimer (**Supplemental Fig. 11**)^20,41^. The E2α’ helix is also sufficient to confer sensitivity to RhoA-mediated activation. Replacement of the PLCβ3 E2α-F2α helices in the EF hands (residues 183-221) with the corresponding region of PLCε (F2α-E2α’, PLCε residues 1196-1284), or insertion of E2α’ between PLCβ3 EF1/2 and EF3/4 subdomains (PLCβ3 residues 221-222), resulted in increased activity upon cotransfection with RhoA^G14V^ **(Supplemental Fig. 12**).

Residues in PLCε that interact with RhoA×GTP in the structure were also mutated to test their role in activation (**Fig. 4c, Supplemental Fig. 7**). PLCε Arg1049, Trp1051, Phe1187 and Val1189 are positioned to interact with the C-terminus of the switch II helix in RhoA (**Fig. 4c**). PLCε R1049A, in the EF1 module, and F1187E and V1189E, in the loop immediately preceding the E2α’ helix, had ∼2-fold lower maximum activity. PLCε W1051E resulted in a significant ∼5-fold reduction in its maximum activity, confirming the importance of this residue in packing against switch II (**Fig. 4c**). PLCε Ile1279, Ala1282, Ile1283, and Ala1286 are located on the surface of the E2α’ helix and interact with Phe39, Leu69, and Leu72 in the switch regions of RhoA·GTP (**Fig. 4c**). PLCε I1279E had a ∼2-fold decrease in its maximum activity, whereas A1282K, I1283E, and A1286K significantly decreased maximum activity, with the I1283E mutation eliminating the response to the G protein (**Fig. 4d**). PLCε Ile1295E, located on the loop connecting E2α’ back to EF3, also caused a significant ∼4-fold decrease in maximum activity relative to wild-type (**Fig. 4d**). Together, PLCε Trp1051, the E2α’ helix, and Ile1295 make the required interactions with the G protein for RhoA-dependent activation.

## DISCUSSION

In this study, we provide the first molecular insights into the mechanism by which RhoA activates PLCε. Using cell-based assays, we show that the EF hands are critical for RhoA-dependent activation, whereas the Y-box, previously thought to be involved in the process, is needed for general lipase function. The role of the Y-box is not yet known, but because these variants are detected by western blot at the correct molecular weight, the lack of activity seems unlikely to reflect a folding defect. To define the regions of the EF hands necessary for activation, we generated chimeras wherein the PLCε EF hands were replaced, in whole or in part, with those of PLCβ3. However, only a chimera containing the PLCβ3 EF1/2 subdomain could be activated by RhoA in cells (**Figs. 1,2**). We then used cryo-EM to determine a reconstruction of the RhoA×GTP–PLCε PH-C complex (**Fig. 3**). Although the quality of the overall RhoA×GTP density was weak, the overall shape of the density was consistent and the switch regions could be resolved, allowing us to fit RhoA in the map. Site-directed mutagenesis and cell-based assays support the model for the complex. RhoA×GTP binds to the E2α’ helix in the PLCε EF hands, making additional contacts with residues in EF1 and the loop connecting E2α’ to the EF3/4 module. Deletion or mutation of the E2α’ helix, EF1/2, or the loop connecting E2α’ and EF3/4 all but eliminates RhoA-dependent activation (**Fig. 4**) but has little impact on activation by other G proteins (**Supplemental Fig. 11)**. These findings strongly indicate that E2α’ is the binding site for RhoA.

PLCε must interact with the inner leaflet of the plasma membrane to hydrolyze its PIP_2_ substrate. In addition to the PIP_2_ binding site in the active site of the TIM barrel, the CDC25 and PH domains contribute to membrane association, and together define a common membrane engagement surface^18^. In the reconstruction of the RhoA×GTP–PLCε PH-C complex, the C-terminal thirteen residues of the GTPase, which are disordered in prior atomic structures, allow the prenylated C-terminus of RhoA to reach the same plane of the predicted membrane surface. Thus, the architecture captured in our reconstruction is compatible with functional complex at a membrane. Although membrane localization is an important component of the RhoA activation mechanism, we showed this alone is insufficient for maximum activation, as soluble, active RhoA proteins stimulate PLCε PH-C to a submaximal threshold in liposome-based assays (**Supplemental Fig. 2**). RR distance plots comparing the structures of PLCε PH-C^18^ and PLCε EF3-RA1^30^ show that the EF3/4 module is closer to the TIM barrel domain when RhoA×GTP is bound (**Supplemental Fig. 8**). This may promote rearrangements within the catalytic domain that, when the complex is at the membrane, facilitate displacement of the autoinhibitory X–Y linker and allow substrate binding (**Fig. 5**). This is supported by mutations in EF3/4 and the TIM barrel that decrease maximum RhoA-dependent activation ∼3-fold (**Supplementary Fig. 10**). The role of the Y-box however remains undetermined but is clearly functionally important because its deletion impairs both basal and G protein-stimulated activities (**Fig. 1a,b**).

**Figure 5.**
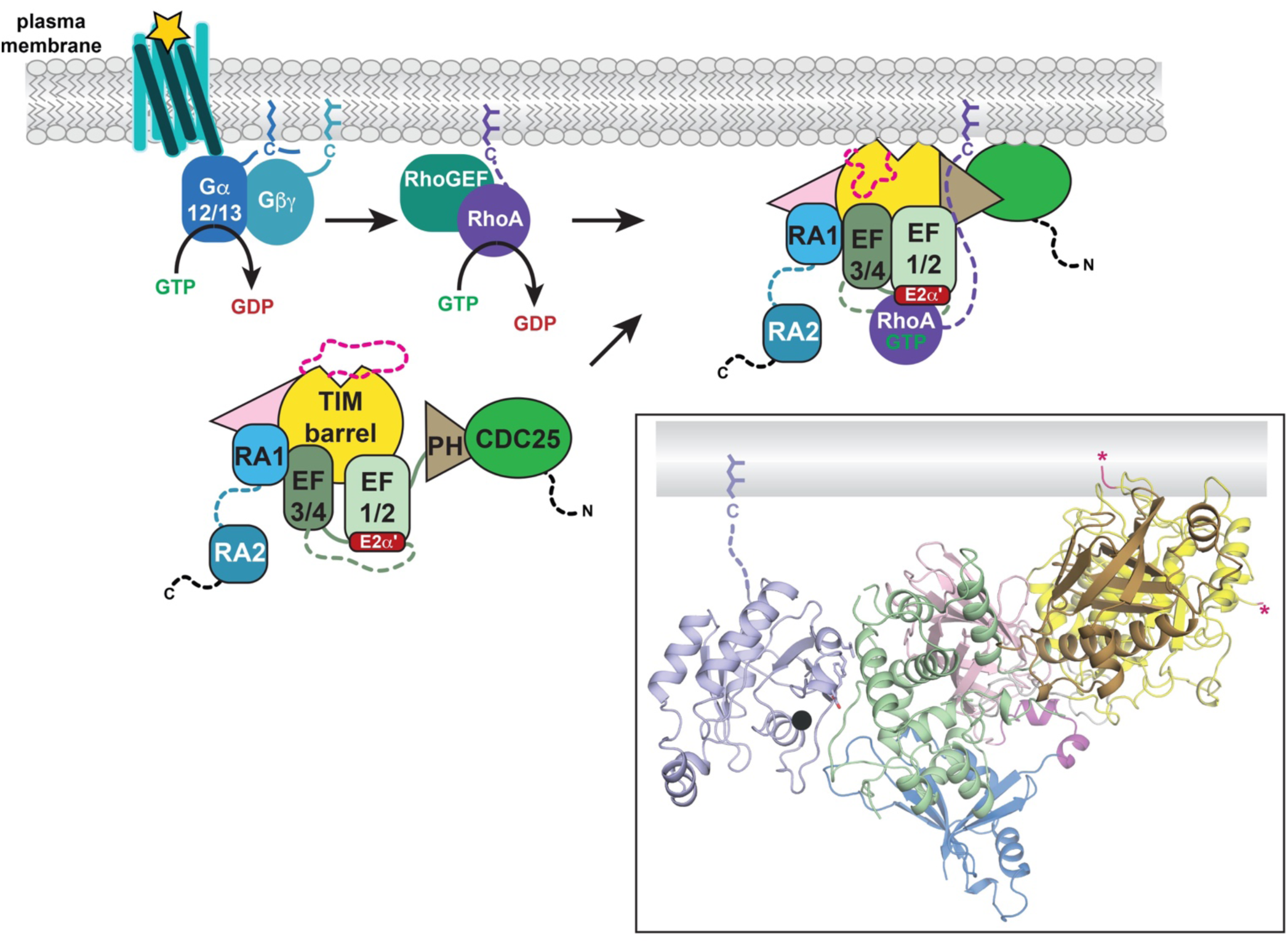
Schematic for RhoA-mediated PLCε activation. Under basal conditions, PLCε is autoinhibited by the X–Y linker in the cytoplasm. RhoA is activated downstream of G_12/13_ coupled receptors by a RhoA guanine nucleotide exchange factor (RhoGEF). The activated G protein binds the lipase via the E2α’ helix, which together with the PLCε CDC25, PH, and TIM barrel domains, defines a common membrane interaction surface. RhoA binding induces conformational changes within the lipase, moving the EF hands closer to the TIM barrel domain, stabilizing the EF1/2-EF3/4 interface and the CDC25-PH domain module closer to the TIM barrel. These long-range conformational changes may facilitate displacement of the X–Y linker from the active site, exposing the active site. In the boxed inset, the RhoA×GTP–PLCε PH-C reconstruction is shown. The prenylated C-tail of RhoA is shown as a dashed line and together with the TIM barrel and PH domain, forming a membrane interaction surface. The ends of the PLCε X–Y linker are indicated by hot pink asterisks.

Whether other G protein activators of PLCε regulate lipase activity through similar mechanisms has yet to be determined. It seems that membrane localization will always be a key component because the autoinhibitory X–Y linker would need to be displaced via interfacial activation at the membrane^3,21^, and all known activators of PLCε are prenylated G proteins^3^. At a minimum, regulation by Gβγ and Rap1A occurs independently of the PLCε E2α’ helix because its deletion did not alter the fold change in activity by these G proteins. The Gβγ binding site has not been determined, but the CDC25 and RA2 domains are both required for maximum activation^41^. It is possible that two Gβγ molecules bind to the lipase, promoting translocation to the membrane, a scenario that has been observed in other Gβγ–effector enzymes complexes, such as the Gβγ–PI3K^42^ and Gβγ–PLCβ3^43^ complexes. In the case of Rap1A, the GTPase binds the C-terminal RA2 domain of PLCε, but activation requires the PH and EF1/2 domains indicating that there are allosteric consequences. Indeed, small-angle X-ray scattering showed that binding of constitutively active Rap1A to PLCε PH-C induced long-range conformational changes, stabilizing the lipase in an extended state^20^. This could potentially be explained either by the Rap1A-RA2 module interacting with the PH and/or EF1/2 domains, or by the latter two domains constituting a second Rap1A binding site^3^.

In the future, studies that integrate the role of the membrane in PLCε function are needed for a complete understanding of regulation. PLCε translocates to the cytoplasmic leaflets of the plasma and perinuclear membranes, hydrolyzing PIP_2_ or PI4P, respectively^2,3^. These membranes differ in their composition and biophysical properties, and whether and how these factors impact interfacial activation, membrane engagement, and/or G protein activation remain to be explored.

## METHODS

### Cloning of PLCε and variants

A pCMV vector encoding *R. norvegicus* PLCε with a C-terminal FLAG tag (gift from A.V. Smrcka, U. Michigan) was used as a template to generate PLCε PH-C (residues 837-2282), EF-C (residues 1038-2282), and EF3-C (residues 1284-2282). The Y-box (residues 1667-1728) was deleted in PLCε and EF-C by Q5 site-directed mutagenesis (New England BioLabs, Inc.). In-Fusion cloning (Takara Bio USA, Inc.) was used to generate PLCε and PLCβ3 (in a pCI-neo vector, Promega) chimeras. For PLCε/β3 EF, PLCε residues 1038-1355 were replaced with PLCβ3 152-304, PLCε/β3 EF1/2 replaced PLCε residues 1038-1284 with residues 152-216 from PLCβ3, and PLCε/β3 EF3/4 replaced PLCε residues 1035-1355 with PLCβ3 residues 217-304. Internal deletions in the PLCε EF hands (Δ1275-1289, Δ1287-1298, and Δ1275-1298) and point mutants were generated with In-Fusion cloning (Takara Bio USA, Inc.). PLCε PH-C was subcloned into pFastBac HTA (ThermoFisher) for protein expression and purification^41^. All constructs were sequenced over the coding region.

### Expression and purification of PLCε variants

*R. norvegicus* PLCε PH-C was expressed in baculovirus-infected *Spodoptera frugiperda* (Sf9) cells at an MOI of ∼1 for 48 h and harvested by centrifugation. The pellets were resuspended in lysis buffer (20 mM HEPES pH 8.0, 50 mM NaCl, 10 mM β-mercaptoethanol (β-Me), 0.1 mM EDTA, 0.1 mM EGTA, and EDTA-free protease inhibitor tablets (Roche) at one-third strength), homogenized and lysed by dounce, and centrifuged at 100,000 x *g* for 1 h. The supernatant was filtered and diluted to a final volume of 320 mL with 20 mM imidazole, 300 mM NaCl, and 10 mM β-Me prior to loading on a 5 mL HisTrap column pre-equilibrated with binding buffer (20 mM HEPES pH 8.0, 300 mM NaCl, 10 mM β-Me, 0.1 mM EDTA, 0.1 mM EGTA, and 20 mM imidazole). The column was washed with 8 column volumes (CVs) of binding buffer and eluted with a 0-500 mM imidazole gradient. Fractions containing the protein were concentrated to ∼1 mL, and exchanged into low salt buffer (20 mM HEPES pH 8.0, 50 mM NaCl, 0.1 mM EDTA, 0.1 mM EGTA, and 2 mM DTT) before loading on a 1 mL MonoQ column pre-equilibrated with the low salt buffer. The protein was eluted with a 0-500 mM NaCl gradient. Fractions containing the protein were pooled, concentrated to ∼1 mL, and then applied to tandem Superdex 200 Increase 10/300 GL columns (Cytiva) equilibrated with 20 mM HEPES pH 8.0, 200 mM NaCl, 0.1 mM EDTA, 0.1 mM EGTA, and 2 mM DTT. Fractions containing the final, purified protein were identified by SDS-PAGE, concentrated to 4-5 mg/mL, flash-frozen in liquid nitrogen, and stored at -80 °C.

### Cloning of RhoA variants

The cDNA encoding wild-type human RhoA (residues 1-194, gift from J.J.G. Tesmer, Purdue U.) was subcloned into pcDNA 3.1 and an HA-tag installed at the N-terminus. Soluble RhoA was generated by subcloning the human cDNA into a pMALc2H_10_T vector (gift from J. J. G. Tesmer, Purdue). The G14V mutation was introduced using Q5-site directed mutagenesis (New England BioLabs, Inc.). RhoA and RhoA^G14V^ were also subcloned into pFastBac HTA for expression and purification. All constructs were sequenced over the coding region.

### Expression and purification of RhoA variants

RhoA and RhoA^G14V^ were expressed in baculovirus-infected High5 cells, cultured in Lonza Insect-XPRESS media (Fisher Scientific) at an MOI of ∼1 and harvested after 48 h. Pellets were resuspended in lysis buffer (20 mM HEPES pH 8.0, 150 mM NaCl, 0.1 mM EDTA, 10 mM β-Me, 10% glycerol, 20 mM GDP, 1 mM leupeptin and lima bean (LL) protease, 1 mM phenylmethanesulfonylfluoride (PMSF), and 5 mM MgCl_2_). Cells were lysed by four freeze-thaw cycles and centrifuged at 100,000 x *g* for 1 hour. The membrane pellet containing RhoA or RhoA^G14V^ was resuspended in solubilization buffer (20 mM HEPES pH 8.0, 150 mM NaCl, 10 mM β-Me, 10% glycerol, 20 mM GDP, 1 mM LL, 1 mM PMSF, and 5 mM MgCl_2_) and sodium cholate added to a final concentration of 1% (w/v). The slurry was stirred for 1 h at 4 °C to solubilize the membrane fraction, then centrifuged at 100,000 x *g* for 45 min. The supernatant was diluted 5-fold with load dilution buffer (solubilization buffer supplemented with 1% (w/v) sodium cholate) and loaded on an Ni-NTA affinity column (Roche cOmplete Ni-NTA resin) equilibrated with 10 CVs load dilution buffer. The column was washed with 10 CV of wash 1 buffer (20 mM HEPES pH 8.0, 150 mM NaCl, 10 mM β-Me, 10% glycerol, 20 mM GDP, 1 mM MgCl_2_, and 1% (w/v) sodium cholate) and 20 CVs of wash 2 buffer (20 mM HEPES pH 8.0, 300 mM NaCl, 10 mM β-Me, 10% glycerol, 20 mM GDP, 1 mM MgCl_2_, 10 mM CHAPS, and 20 mM imidazole). RhoA or RhoA^G14V^ was eluted in 10 CVs of elution buffer (20 mM HEPES pH 8.0, 150 mM NaCl, 10 mM β-Me, 10% glycerol, 20 mM GDP, 1 mM MgCl_2_, 10 mM CHAPS, and 250 mM imidazole), and concentrated to ∼1 mL. The protein was loaded on a Superdex 75 Increase 10/300 GL (Cytiva) column equilibrated with 20 mM HEPES pH 8.0, 150 mM NaCl, 1 mM DTT, 40 mM GDP, 1 mM MgCl_2_, and 6 mM CHAPS. Fractions containing the purified protein were identified by SDS-PAGE, flash frozen in liquid nitrogen, and stored at -80 °C.

For soluble RhoA and RhoA^G14V^, *E. coli* BL21(DE3) were transformed and grown to OD_600_ of 0.4-0.6. Expression was induced by addition of 1 mM isopropyl β-*D*-1-thiogalactopyranoside (IPTG) for 16-18 h at 18 °C. Cells were harvested by centrifugation and flash frozen in liquid nitrogen. Proteins were purified as described above, with some modifications. Sodium cholate and CHAPS were omitted from all buffers. After the pellets were resuspended in lysis buffer, 1 mg/mL lysozyme (Fisher BioReagents) was added and incubated on ice for 30 min. To ensure complete lysis, samples were then sonicated on ice for 20 cycles (15 sec pule, 45 sec recovery), then centrifuged at 100,000 x *g* for 1 h. After Ni-NTA affinity chromatography, the protein concentration was measured using a Bradford assay and incubated with 8% (w/w) TEV protease overnight at 4 °C in dialysis buffer (20 mM HEPES pH 8.0, 150 mM NaCl, 10 mM β-Me, 10% glycerol, 40 mM GDP, and 5 mM MgCl_2_). The next day, the Ni-NTA column was washed with 20 CV of dialysis buffer, and the dialysate was passed five times over the resin. The TEV-cleaved, soluble RhoA or RhoA^G14V^ was concentrated to ∼1 mL, and purified on tandem Superdex 75 Increase 10/300 GL (Cytiva) columns.

Purified RhoA proteins were activated using nucleotide exchange. Briefly, the proteins were incubated with a 10-fold molar excess of GTP or GTPγS and a 4-fold molar excess of EDTA for 1.5 h on ice. The reaction was quenched by addition of a 10-fold molar excess of MgCl_2_ and incubatied for 30 min on ice^23^. The proteins were flash frozen in liquid nitrogen and stored at -80 °C.

### Rap1A^Q63E^ and Avi-Gβγ cloning

The cDNA encoding human Rap1A (residues 1-184) was subcloned into pcDNA3.1 with an N-terminal HA tag. The Q63E mutation was introduced using the Q5-Site Directed Mutagenesis Kit (New England BioLabs Inc). Human Avi-tagged Gβ1 and Gγ2 were subcloned into a pCI-neo vector (gift from A.V. Smrcka, U. Michigan). Constructs were sequenced using whole-plasmid sequencing (Plasmidsaurus).

### [^3^H]-IP_x_ accumulation assay

COS-7 cells (gift from A.V. Smrcka, U. Michigan) were seeded at a density of 100,000 cells/well in 12-well plates in Dulbecco’s Modified Eagle’s Medium (Corning) supplemented with 10% fetal bovine serum (FBS, BioTechne), 1% Glutamax (Gibco), and 1% penicillin-streptomycin (Corning) and incubated for 24 h. The cells were co-transfected with 750 ng empty pCMV, 750 ng PLCε variant DNA alone or with 375 ng of RhoA^G14V^, 375 ng of Rap1A^Q63E^, or 375 ng Avi-Gβ1 and 375 ng Gγ2 DNA. After 24 h, the cells were washed with serum- and inositol-free Ham’s F-10 media (Invitrogen),and incubated for 16-18 hours in Ham’s F-10 media supplemented with 1.5 mCi/well myo[2-^3^H(N)] inositol (Revvity). 10 mM LiCl was added to each well and incubated for 1 h to inhibit inositol phosphatases. The media was aspirated, cells were washed once with ice-cold PBS. Cells were lysed on ice by addition of 1 mL ice-cold 50 mM formic acid. Lysates containing [^3^H]-labelled inositol phosphates were loaded onto pre-equilibrated Dowex AGX8 anion exchange columns (BioRad), washed twice with 50 mM formic acid, once with 100 mM formic acid, eluted with 1.2 M ammonium formate and 0.1 M formic acid into scintillation vials. Total [^3^H]-IP_x_ was quantified by scintillation counting (Uniscint BD scintillation cocktail, National Diagnostics)^29,30^. All experiments were performed at least three times in triplicate from independent transfections.

### Immunoblotting

Cells were plated, transfected, and incubated in Ham’s F-10 media 24 h post-transfection, replicating the conditions used for the [^3^H]-IP_x_ accumulation assays. After 48 h, cells were washed once with cold PBS and scraped into 100 μL of 1X SDS loading dye (100 mM Tris-HCl pH 6.8, 6% w/v sucrose, 2% w/v SDS, 5% v/v β-Me, and 0.02% bromophenol blue) and incubated at 90 °C for 10 min prior to loading on a 10% SDS-PAGE gel. Samples were transferred to a PVDF membrane overnight in Towbin buffer (25 mM Tris, 192 mM glycine, 20% (v/v) methanol) at 4 °C. The membrane was blocked with 5% BSA in 1X Tris-buffered saline supplemented with 0.1% Tween-20 (TBST) for 1 h, washed three times with 1X TBST, and incubated overnight at 4 °C with an anti-FLAG rabbit antibody, anti-HA rabbit or mouse antibody, and anti-actin mouse antibody (Cell Signaling Technology) at 1:1000 dilutions. Endogenous and transfected Avi-Gβγ were detected using an anti-GNB1 rabbit antibody (Invitrogen). The next day, the blot was washed three times in 1X TBST and incubated with goat anti-mouse or anti-rabbit secondary antibody conjugated with HRP (Sigma Aldrich) at a 1:10,000 dilution at 27 °C for 1 h. The blot was washed three times in 1X PBS and the West Pico ECL substrate was added (ThermoFisher Scientific)^30^. Blots were imaged using a GeneGnome and densitometry analysis performed in ImageJ^24^.

### Liposome-based activity assays

100 μM of hen egg white phosphatidylethanolamine (PE) and 250 μM of soybean phosphatidylinositol (PI, Avanti) were mixed, dried under nitrogen, and stored at -20 °C. Lipids were resuspended by bath sonication in buffer containing 50 mM HEPES pH 7.4, 80 mM KCl, 2 mM EGTA, and 1 mM DTT. Purified PLCε PH-C was diluted to a final amount of 2-5 ng in assay buffer (100 mM HEPES pH 7.4, 160 mM KCl, 6 mM EGTA, and 1 mM DTT), 3 mg/mL bovine serum albumin (BSA), and 3 mM DTT). RhoA buffer (20 mM HEPES pH 8, 150 mM NaCl, 1 mM DTT, 40 mM GTP, 1 mM MgCl_2_, and 6 mM CHAPS) alone or containing a final concentration of 3 mM RhoA variant was then added. All samples were transferred to 30 °C and incubated with 10 μL liposomes for 2 min before the reaction was initiated by addition of 5 μL free Ca^2+^ solution (1X assay buffer, 1 mM DTT, and 18 mM CaCl_2_). Reactions were incubated for 15 min at 30 °C, then quenched with 5 μL Ca^2+^ chelating solution (1X assay buffer, 1 mM DTT, and 210 mM EGTA). Negative controls lacked free Ca^2+18^. All assays were performed at least three times in triplicate using proteins purified from at least two independent preparations.

### Cryo-EM Sample Preparation

PLCε PH-C at 0.6 mg/mL was incubated with wild-type RhoA•GTP in a 1:3 molar ratio in 20 mM HEPES pH 8.0, 150 mM NaCl, 2 mM DTT, 0.1 mM EDTA, 0.1 mM EGTA, 1 mM MgCl_2_, 40 mM GTP, 0.5 mM CaCl_2_ and incubated on ice for 1 h. The reaction was supplemented with CHAPS (Millipore Sigma) to a final concentration of 2.5 mM and 3.5 μL of the reaction was applied to a glow-discharged Quantifoil 1.2/1.3 300 mesh holey copper grid. Grids were blotted at blot force 2 for 3 seconds, at 4.2 °C with 100% humidity, and plunge frozen in liquid ethane using a Vitrobot Mark IV (ThermoFisher Scientific).

### Cryo-EM data acquisition

Grids containing RhoA•GTP–PLCε PH-C were imaged on a Titan Krios G4 (ThermoFisher Scientific) electron microscope equipped with a post-GIF K3 Summit Direct Electron Detector (Gatan, Inc.) and Gatan quantum GIF energy filter. 6,378 movies were collected using 300 kV at a magnification of 81,000 X (pixel size of 0.527 Å) and defocus range of 0.6-2.0 μm using EPU^44^. Each movie stack recorded 40 frames, for a total dose of 57.8 electrons/Å^2^, and a total exposure time of 3.21 seconds per stack.

### Cryo-EM data processing

The cryo-EM workflow and validation for the RhoAߣGTP–PLCε PH-C complex are shown in Supplemental Figs. 4, 5 and Supplemental Table 1. Patch motion correction and contrast transfer function (CTF) were calculated using cryoSPARC^45^. From a total of 6,378 exposures, 1,329,298 particles were accepted. As a first step, ∼400 particles were manually selected, generating twenty 2D classes for template picking, and fifty 2D class averages generated. Nine classes (291,245 particles) were selected for an *ab initio* reconstruction that generated two initial models, Volumes A1 and B2. Volume A1 was larger and contained more particles, and so was used as the input for a non-uniform refinement to generate Volume A2 (3.52 Å, 184,875 particles). An AlphaFold2 model of PLCε PH-C, as well as the PLCε PH-C reconstruction from a Fab-bound complex (PDB ID 9B13^18^) could be fit in the density of Volume A2. Additional, unmodeled density was observed adjacent to the EF hands, and heterogenous refinement was carried using the particles in Volumes A2 and B1, as well as 291,245 particles identified in the *ab initio* reconstruction, to improve the resolution of this region. The resulting Volume A3 (194,157 particles) was subjected to non-uniform refinement against the same volume class, yielding a 3.44 Å map. Rigid body fitting the PLCε PH-C structure into Volume 4A revealed stronger density adjacent to the EF hands. The crystal structure of RhoA (PDB ID 1S1C^36^) was fit in this density with its switch regions poised to interact with the E2α’ helix. This model was then used to generate twenty 2D volume classes in EMAN, which were used as templates for particle picking from the initial micrographs. 5,828,020 particles were extracted, with 1,047,728 particles accepted after inspection and used to generate fifty 2D classes. The best ten 2D classes (209,103 particles) were selected and used to generate two volumes C1 and D1. Volume C1 was unique compared to the initial volumes (Volumes A1 and B)1, while Volume D1 was very similar to Volume A1. To maximize the particle number for the final reconstruction, the 114,829 particles in Volume C1 were used in a non-uniform refinement, generating Volume C2 at 6.67 Å. These 209,103 particles were combined with the original 291,245 particles, and after removal of duplicates, the remaining 419,418 particles, were used in heterogeneous refinement against volumes A2, B1, and C2. The resulting Volume A5 (209,463 particles) was subjected to non-uniform refinement, resulting in the final 3.32 Å map.

### Model building, refinement, and validation

An AlphaFold2 model of PLCε PH-C and the crystal structure of RhoA·GMPPNP (PDB ID: 1S1C^28^) were rigid body fit into the cryo-EM density using COOT^46,47^. Alternating rounds of manual model building in COOT and refinement in PHENIX^46^ were carried out, guided by the DAQ collaboratory^48^ Stereochemistry of the final model was evaluated using MolProbity and CaBLAM in PHENIX^46,49^. Coordinates for the RhoA•GTP–PLCε PH-C reconstruction, volume map, half maps A and B, refinement mask, and FSC curve for Volume A were deposited in the EMDB and PDB as accession numbers EMD-43927 and 9AX5, respectively. Raw micrograph data was also deposited in EMPIAR-12069.

## ACKNOWLEDGEMENTS

We thank Dr. A.V. Smrcka (U. Michigan) for DNA encoding *R. norvegicus* PLCε, human Avi-tagged Gβ and Gγ, pCI neo vector and COS-7 cells, and Dr. J.J.G. Tesmer for DNA encoding human RhoA and the pMALc2H_10_T vector. We are grateful to Dr. Thomas Klose, Dr. Frank Vago, and Steve Wilson for their guidance in cryo-EM data collection and computational support. This work is supported by NIH 1R01HL141076-01 (A.M.L.) and support from the Purdue Institute for Cancer Research Shared Resource Award to the Purdue Cryo-EM Facility is gratefully acknowledged (P30CA023168). The content is solely the responsibility of the authors and does not necessarily represent the official views of the National Heart, Lung, and Blood Institute or the National Institutes of Health.

## AUTHOR CONTRIBUTIONS

V.O. and A.M.L. designed the experimental approach. V.O., K.S., K.M., E.E.G., and A.M.L. cloned, expressed, and purified PLCε variants. V.O., K.S., K.M., E.E.G.-K. and W.C.H., performed and analyzed activity assays, and V.O. and B.D. carried out western blots. V.O. prepared cryo-EM samples, and with K.S., E.E.G., and I.J.F., collected cryo-EM data. V.O. and A.M.L. modeled the structure. V.O. and A.M.L. wrote the manuscript.

## COMPETING INTERESTS

The authors declare no competing interests.

## CORRESPONDENCE AND MATERIALS

Correspondence and requests for materials should be addressed to Angeline M. Lyon.

## SUPPORTING INFORMATION FOR

**Supplemental Figure 1.**
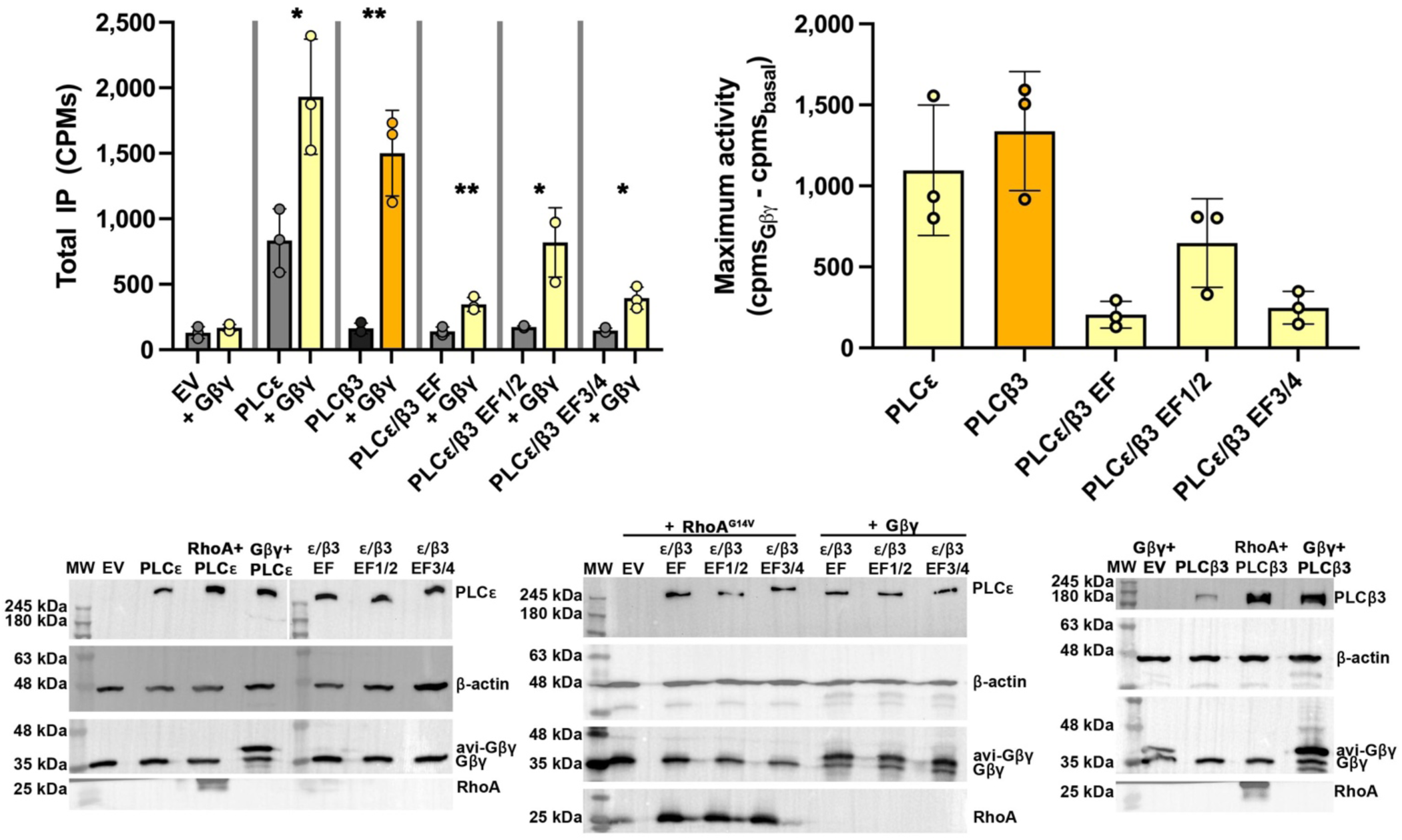
Basal and Gβγ-stimulated activities of PLCε^41^, PLCβ3^22^, and the EF hand chimeras. (*Left*) Data represents three independent experiments from independent transfections in triplicate ± SD and analyzed using unpaired, one-tailed t-test with Welch’s correction to compare the basal and Gβγ-stimulated activities of each variant. For PLCε, *p<0.015; PLCβ3, **<0.0091; PLCεβ3 EF **p<0.0041; PLCε/β3 EF1/2, *p<0.0256, and PLCε/β3 EF3/4, *p<0.0155. (*Right*) The change in maximal activity ± SD was calculated by subtracting RhoA-stimulated activity by the basal activity of each variant. Data was analyzed using a one-way ANOVA and Kruskal-Wallis test comparing each variant to PLCε, followed by a Dunn’s multiple comparisons test. Shown below are representative western blots of PLCε/β3 chimeras co-expressed with either HA-RhoA^G14V^ or Avi-Gβγ, in COS-7 cells for 48 h. Empty pCMV vector (EV) was used as a negative control and β-actin as a loading control. PLCε and PLCβ3 are FLAG-tagged and detected with an anti-FLAG antibody. Endogenous Gβγ and Avi-Gβγ were detected using an antibody that recognizes the Gβ subunit. N-terminally HA-tagged RhoA was detected using an anti-HA antibody.

**Supplemental Figure 2.**
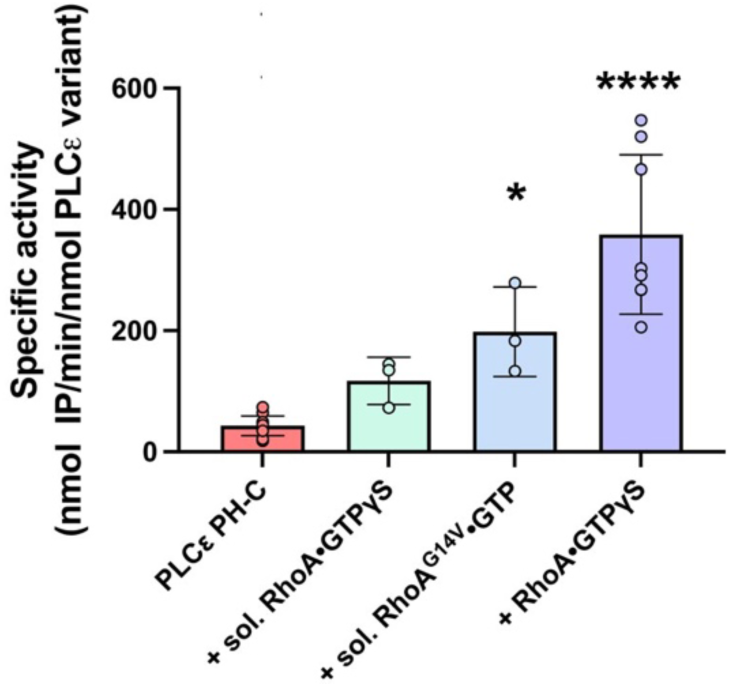
Soluble, active RhoA variants increase lipase activity. Soluble RhoA·GTPγS and RhoA^G14V^·GTP increase the activity of PLCε PH-C to a submaximal threshold in a liposome-based activity assay. Prenylated, wild-type RhoA·GTPγS maximally activates PLCε PH-C. Data shown represents the average ± SD of at least three independent experiments, performed in triplicate, using proteins purified from at two independent purifications. ± SD. Data was analyzed using a one-way ANOVA analysis followed by Dunnett’s multiple comparisons to PLCε PH-C basal activity. *p<0.0221, ****p<0.0001.

**Supplemental Figure 3.**
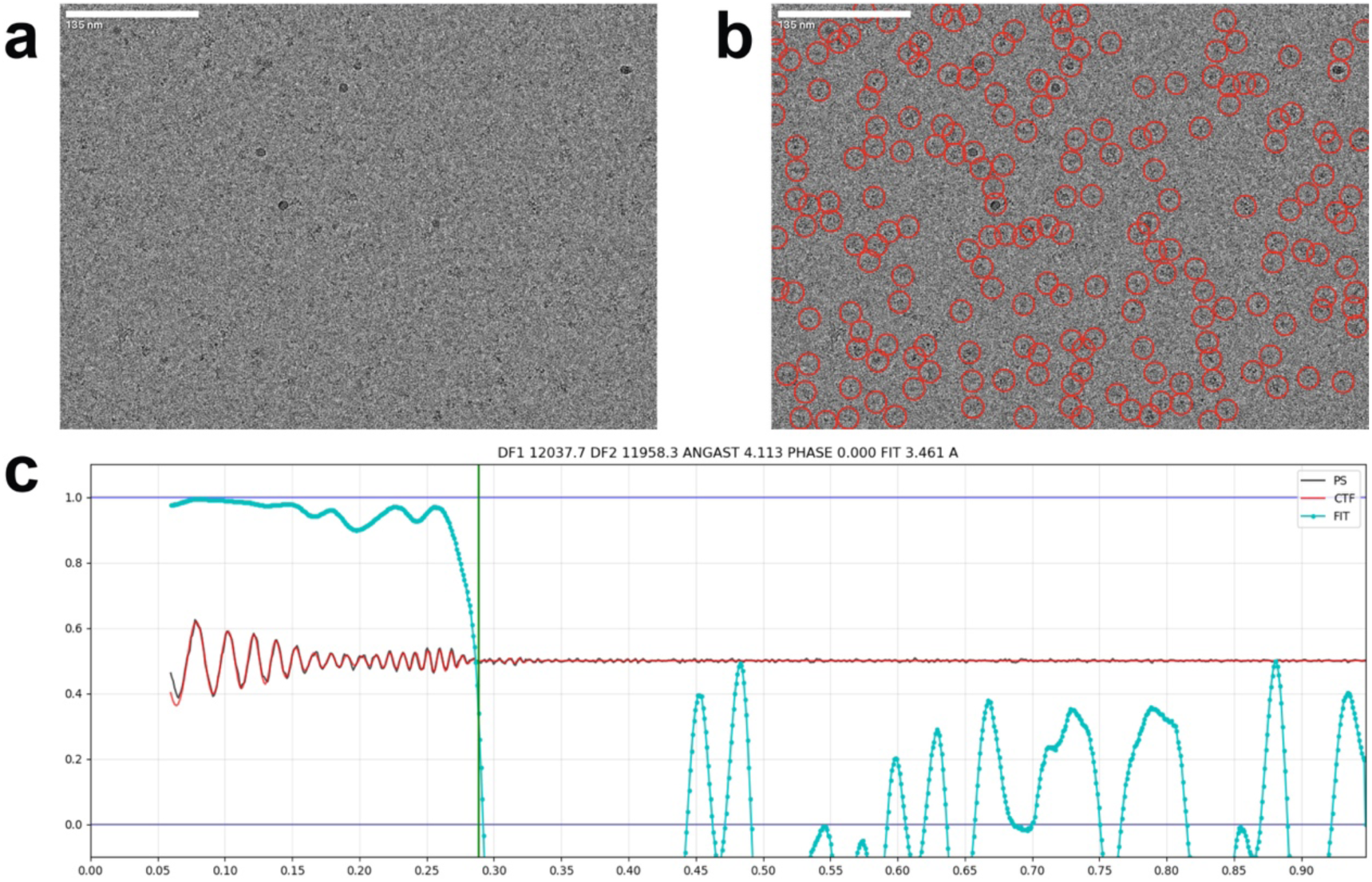
Representative micrograph and CTF plot for the RhoA×GTP–PLCε PH-C complex. (**a**) Representative raw micrograph alone and (**b**) showing template-picked particles circled in red. (**c**) CryoSPARC patch-based 1D CTF (contrast transfer function) fit plot for the micrograph shown in A; x-axis units are 1/Å. Red and black lines show the fit between theoretical (red) and observed (black) Thon rings. The cyan line shows the cross-correlation fit and drops below the 0.5 high-confidence threshold at 3.46 Å.

**Supplemental Figure 4.**
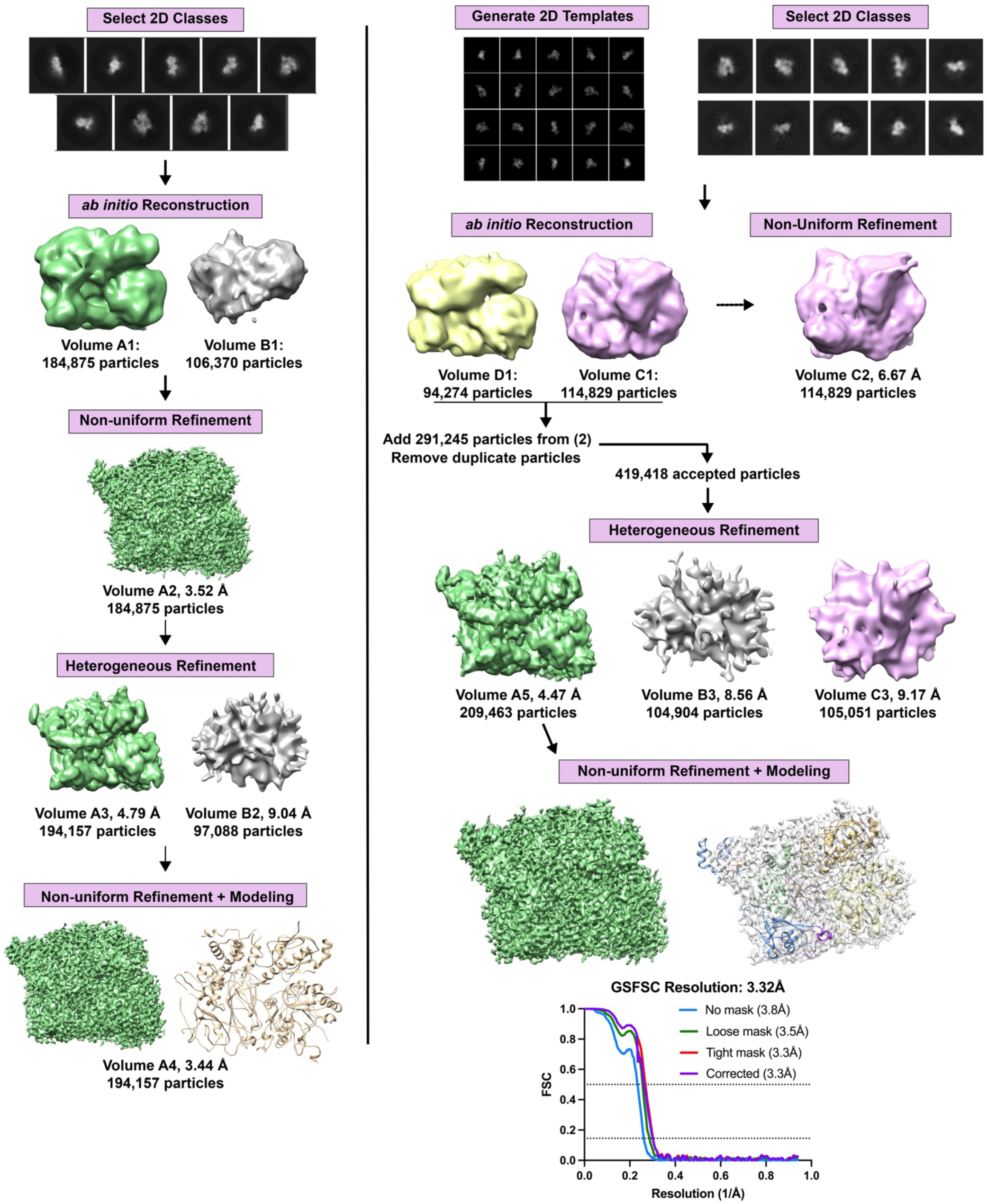
Cryo-EM workflow for the RhoA·GTP–PLCε PH-C reconstruction.

**Supplemental Figure 5.**
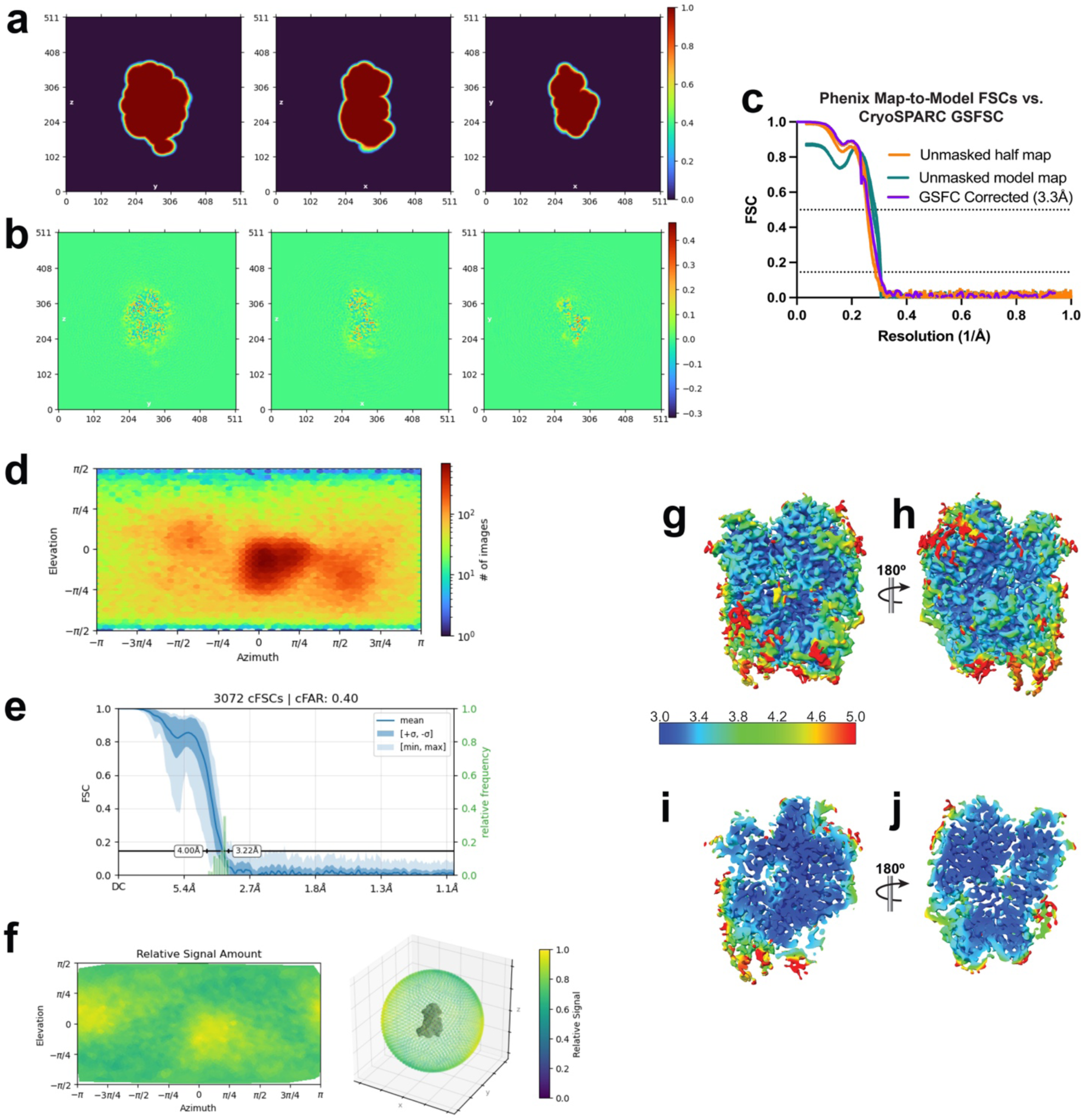
Validation of the RhoA×GTP–PLCε PH-C reconstruction. Final refinement (**a**) real-space mask slices and (**b**) real-space slices. (**c**) FSC curves for the cryoSPARC final 3D map (purple), Phenix map-to-model (teal), and half-map to model (orange). (**d**) CryoSPARC viewing direction distribution plot (Euler angles) of the particles in the structure showing diverse viewing directions, as supported by the (**e, f**) orientation diagnostics plots. The (**e**) conical FSC area ratio (cFAR) plot with a ratio of 0.4 (<0.5) indicates slight preferred orientation. However, the (**f**) relative signal plot and 3DFSC show that most views are well-represented in the dataset. (**g-j**) The refined structure colored by local resolution as calculated in cryoSPARC, including clipped slice views (**i-j**) highlight high-resolution in core PLCε domains.

**Supplementary Table 1.** Cryo-EM data collection, refinement, and validation statistics EMDB: EMD-43927, EMD-43928.

|  |  |
| --- | --- |
| **EMDB:** EMD-43927, EMD-43928 |  |
| **PDB:** 9AX5 |  |
| **EMPIAR:** EMPIAR-12069 |  |
| **Data Collection** |  |
| Grids | Copper Quantifoil |
| Vitrification method | FEI Vitrobot |
| Microscope | FEI Titan Krios |
| Magnification | 81000 |
| Voltage (kV) | 300 |
| Detector | GATAN K3 (6k x 4k) |
| Electron exposure (e-/Å<sup>2</sup>) | 57.8 |
| Number of frames | 40 |
| Defocus range (mM) | 0.6-2.0 |
| Pixel Size (Å) | 0.527 |
| **Data Processing** |  |
| Number of micrographs | 6,378 |
| Initial particle images (no.) | 1,329,298 |
| Final particle images (no.) | 209,463 |
| Symmetry | C1 |
| Map resolution (Å) | 3.3 |
| FSC threshold | 0.143 |
| **Refinement** |  |
| Map sharpening *B* factor (Å<sup>2</sup>) | 133.2 |
| Map CC | 0.80 |
| Model composition |  |
| Non-hydrogen atoms | 8544 |
| Protein residues | 1065 |
| Ligands | 3 |
| *B* factor (Å<sup>2</sup>; min/max/mean) |  |
| Protein | 20.13/162.65/57.21 |
| Ligand | 27.98/138.95/134.91 |
| R.m.s. deviations |  |
| Bond lengths (Å) | 0.004 |
| Bond angles (°) | 0.924 |
| Validation |  |
| MolProbity score | 1.60 |
| Clashscore | 6.17 |
| Rotamer outliers (%) | 1.05 |
| CaBLAM outliers (%) | 1.45 |
| Ramachandran plot |  |
| Favored (%) | 96 |
| Allowed (%) | 4 |
| Outliers (%) | 0 |

**Supplemental Figure 6.**
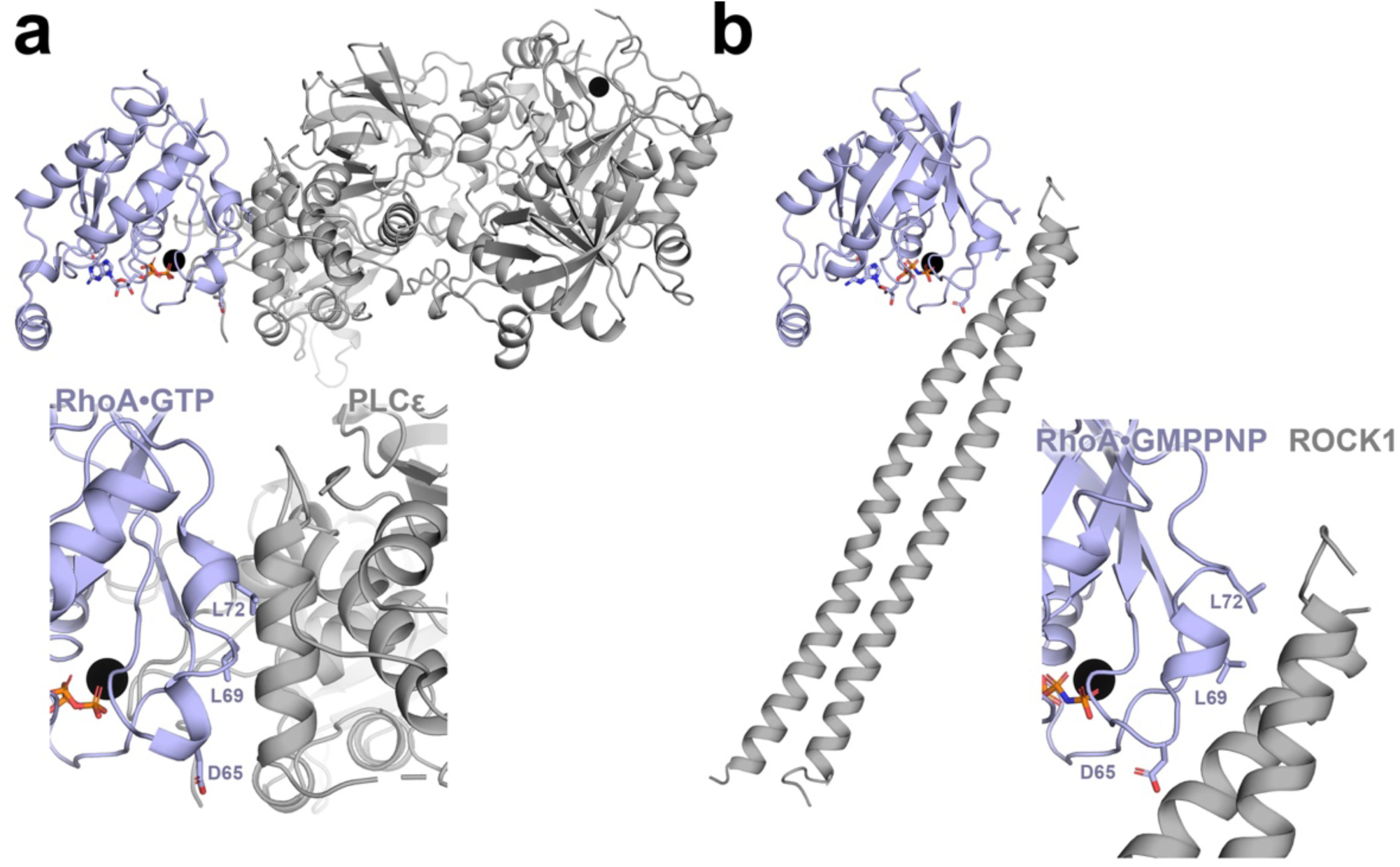
Comparison of RhoA intermolecular interfaces. Active RhoA interacts with **(a)** PLCε PH-C and the **(b)** Rho binding domain (RBD) of Rho-associated coiled-coil containing protein kinase 1 (ROCK1) through similar effector binding sites (PDB ID 1S1C)^36^. The RhoA–PLCε PH-C interface buries ∼1,770 Å^2^ and the RhoA–ROCK1 interface buries ∼1,370 Å^2^ surface area.

**Supplemental Figure 7.**
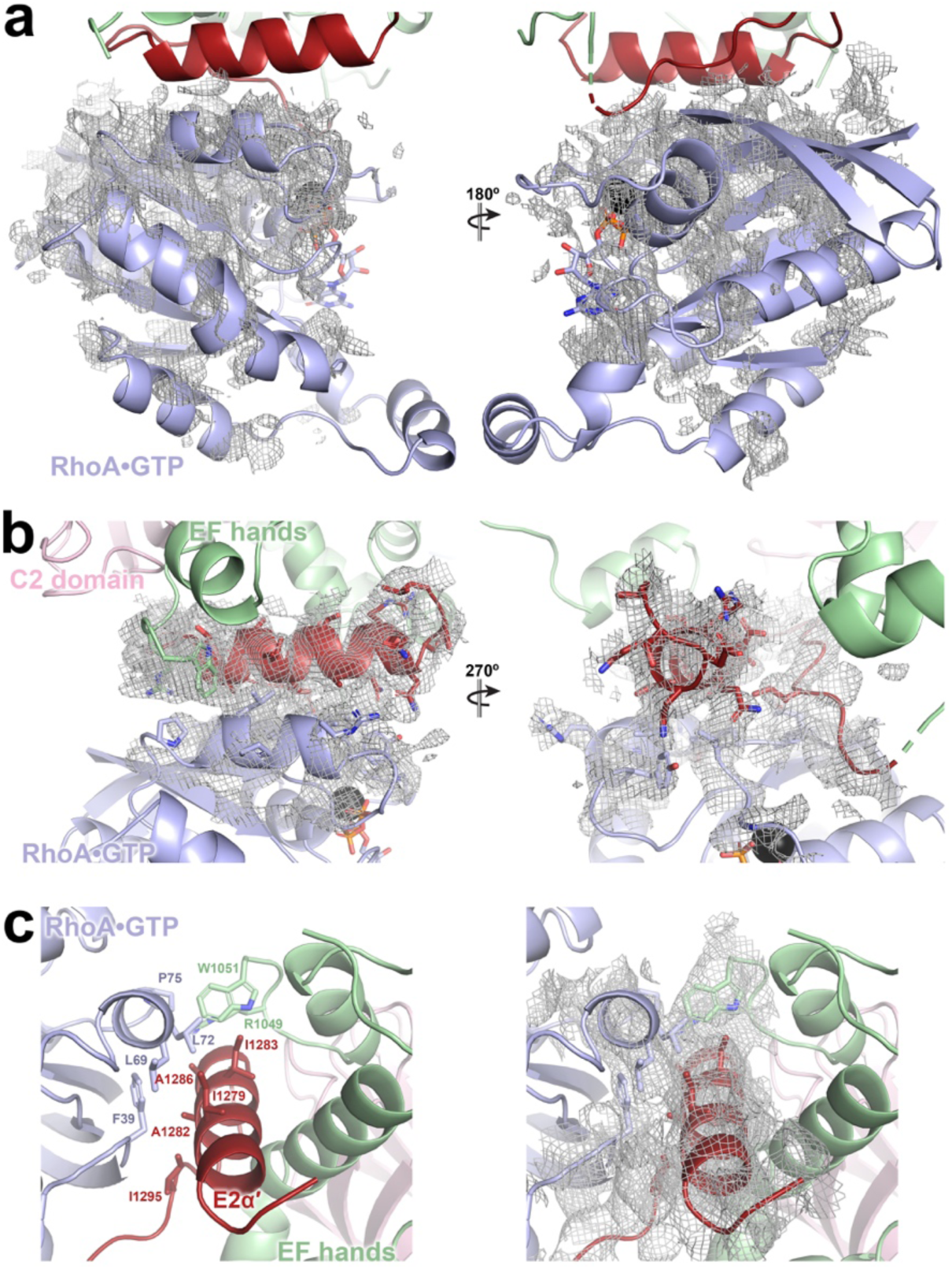
Example cryo-EM density for RhoA×GTP and RhoA×GTP–PLCε PH-C interface. (**a**) Density contoured to 4 α with a 2 Å radius around (**a**) the fitted RhoA**×**GTP molecule and the (**b**) switch regions of RhoA×GTP and the PLCε E2α’ helix and loop, shown in red. (**c**) Residues in the RhoA×GTP–PLCε PH-C interface (as in Fig. 4c) and the corresponding density within the region.

**Supplemental Figure 8.**
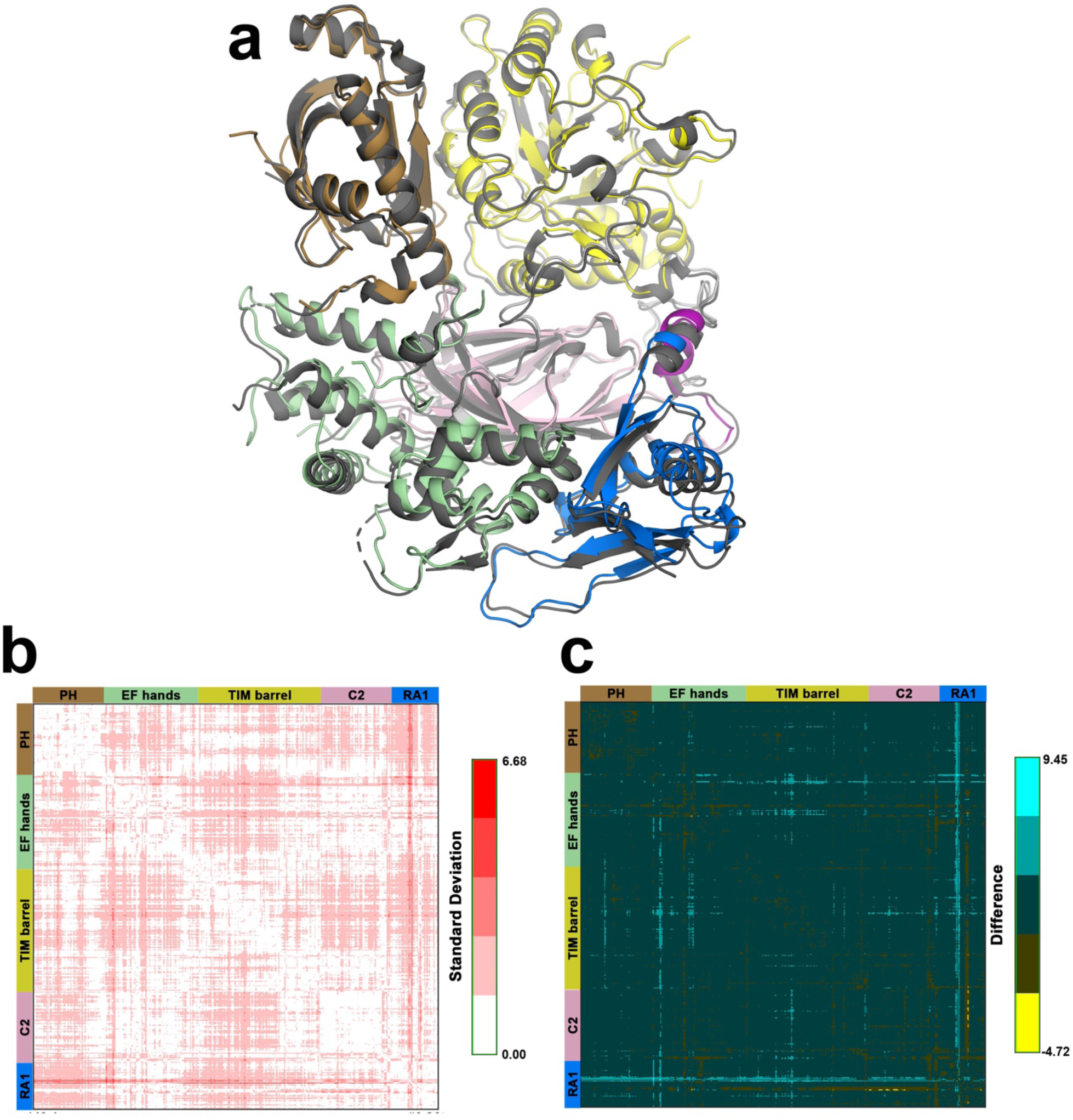
Comparison of PLCε PH-C reconstructions. **(a)** Superposition of the PLCε PH-RA1 domains from the RhoA·GTP–PLCε PH-C reconstruction (colored as in Fig. 1A) and the Fab2– PLCε PH-C reconstructions (dark gray). RhoA·GTP and Fab2 are not shown for clarity. The structures were superimposed over the Y-half of the TIM barrel (residues 1644-1824) based on the RR analysis^40^. The greatest changes are observed in EF3/4 and the RA1 domain. RR distance maps comparing the structures of PH-C when bound to Fab2 or RhoA·GTP (PDB ID 9B13). (**b**) Cα-Cα distances between equivalent residues in the two structures. (**c**) Cα-Cα differences in distance between equivalent residues in the two structures.

**Supplemental Figure 9.**
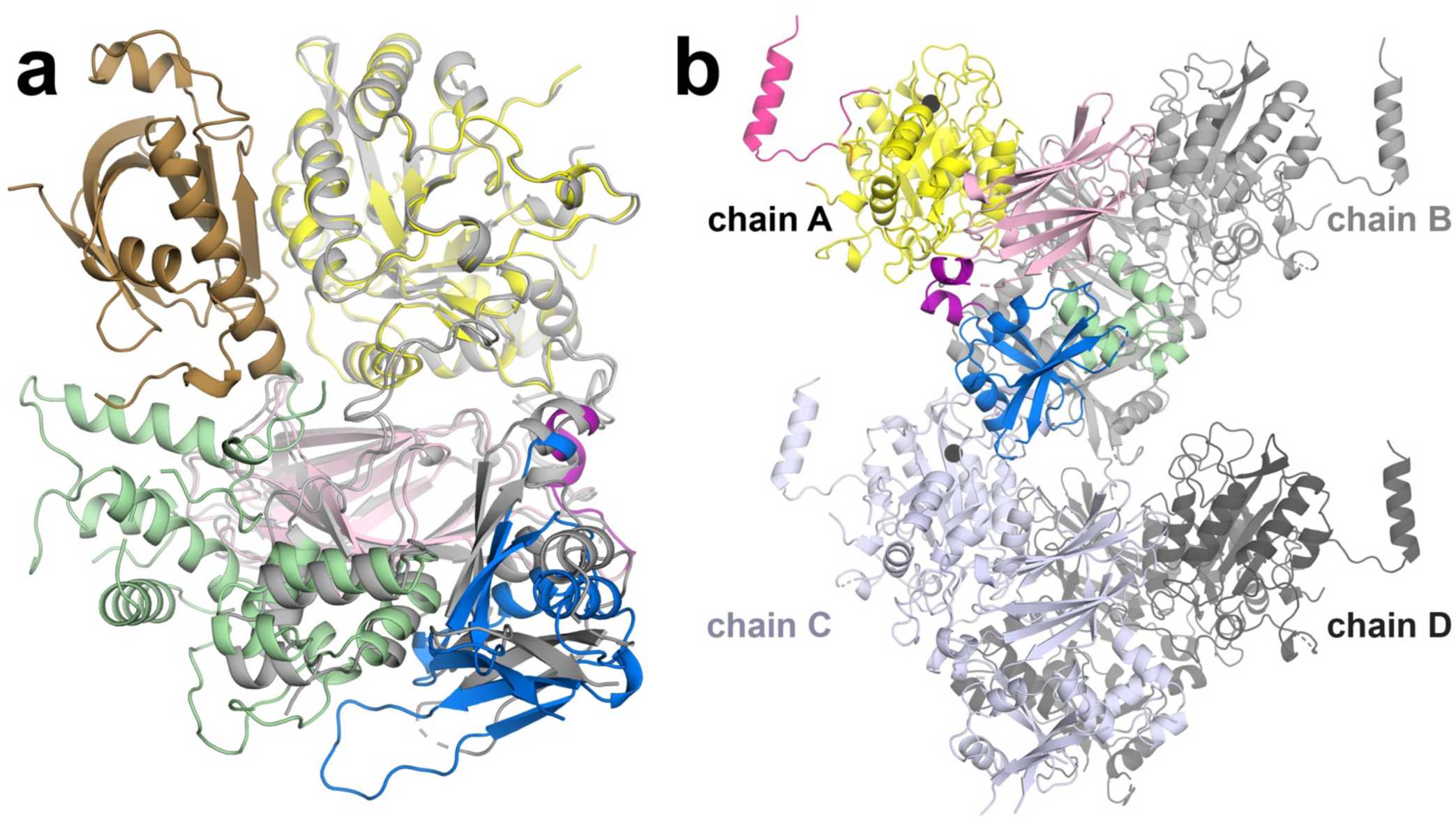
Comparison of PLCε PH-C to the EF3-RA1 crystal structure. (**a**) Superposition of the RhoA·GTP–PLCε PH-C complex with the crystal structure of PLCε EF3-RA1 (PDB ID: 6PMP)^30^, colored as in Fig. 1A and aligned over the Y-half of the TIM barrel. The greatest differences are observed in the EF3/4 subdomain and RA1 domain. These are most likely due to differences in the PLCε construct used for the experiment and lattice interactions within the PLCε EF3-RA1 crystal structure. (**b**) The asymmetric unit of the PLCε EF3-RA1 structure, in which the EF3/4 and RA1 domains contribute to packing interactions. Chain A is colored as in Fig. 1A.

**Supplemental Figure 10.**
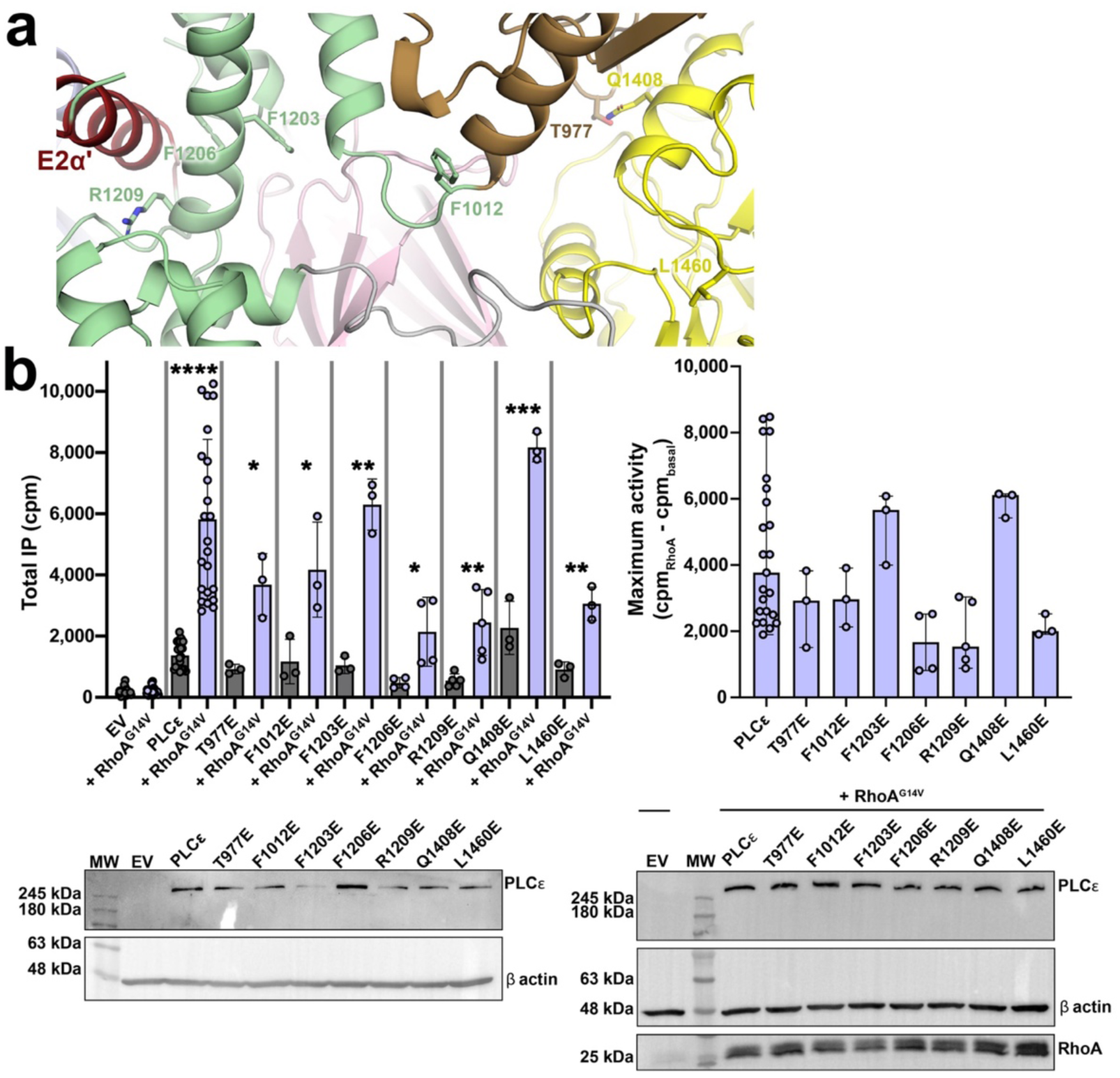
The PLCε F2α helix may relay RhoA binding to the catalytic domain. **(a)** Binding of RhoA·GTP to the E2α’ helix in the EF hands must be relayed to the active site in the TIM barrel domain to increase activity. This could occur through stabilization of the PH domain-TIM barrel interface or the EF3/4-TIM barrel interface. PLCε residues Thr977, Phe1012, Phe1206, and Arg1209 are conserved in PLCε enzymes, while Phe1203, Gln1048, and Leu1460 are conserved across PLC enzymes. **(b)** *Left.* Mutation of residues at domain interfaces had little change on basal activity but decreased RhoA-dependent activation in cells. At least three independent experiments from independent transfections were performed for each variant, and data is shown as the average of triplicate measurements ± SD. Data was analyzed using an unpaired, one-tailed t-test with Welch’s correction comparing the basal and RhoA-stimulated activities of each variant. ****p<0.0001, for T977E *p<0.0199, F1012E *p<0.0303, F1203 **p<0.0023, F1206 *p<0.0288, R1209 **p<0.0078, Q1408E ***p<0.0008, and L14460E **p<0.0052. (*Right*) The F1206E and R1209E mutations, located on the F2α helix, have ∼2-fold lower maximum activity relative to PLCε. T977E in the PH domain, F1012E at the start of EF1/2, and L1460E in the TIM barrel domain also have decreased maximum activity. *Right.* Changes in maximal activity of each variant were analyzed using a one-way ANOVA and Kruskal-Wallis test comparing each variant to PLCε, followed by a Dunn’s multiple comparisons test. Representative western blots are shown below, with empty pCMV vector (EV) and β-actin used as loading controls. PLCε variants express a C-terminal FLAG tag and are detected using an anti-FLAG antibody, while RhoA contains an N-terminal HA tag, and is detected using an anti-HA antibody.

**Supplemental Figure 11.**
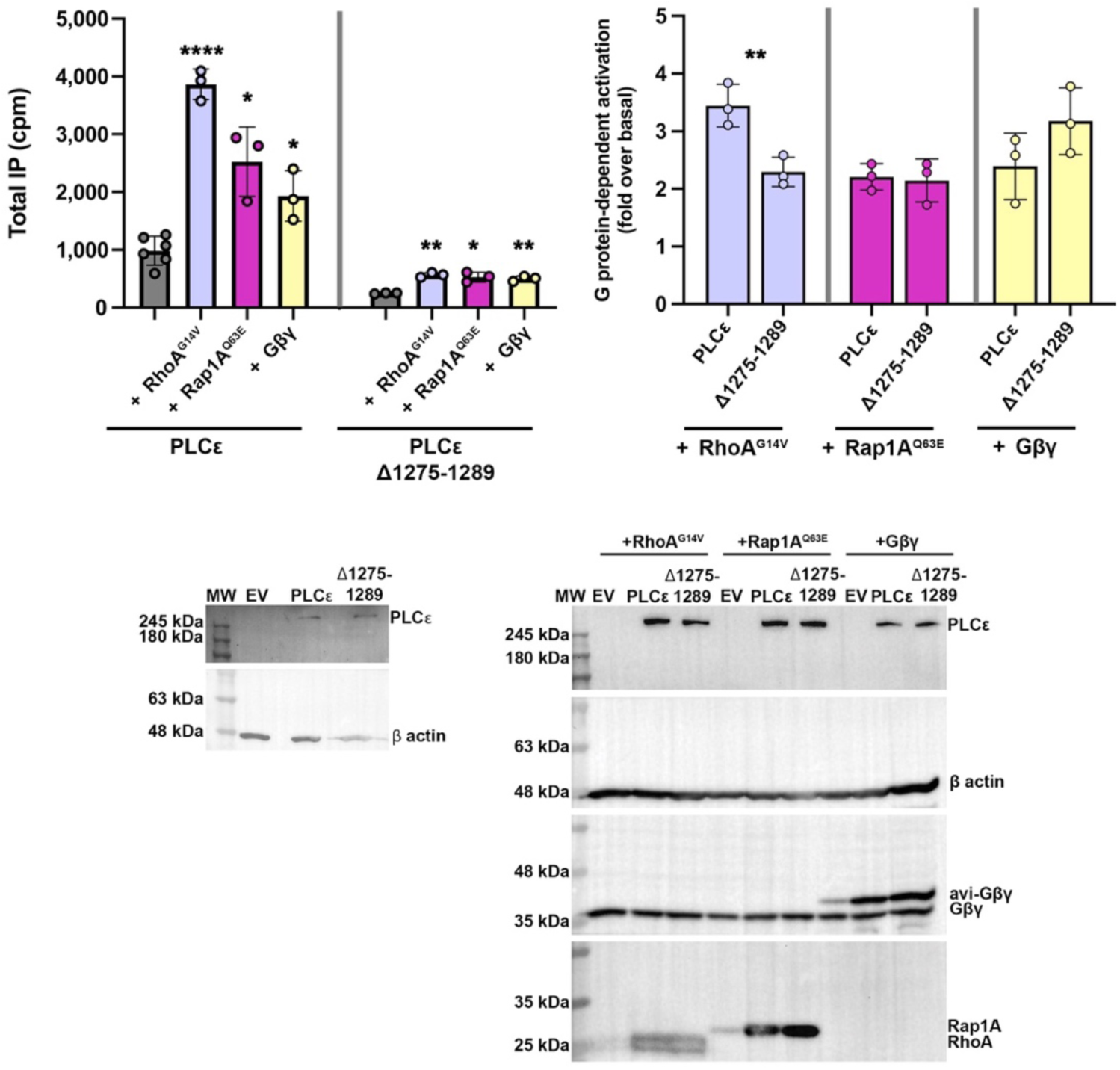
Deletion of the PLCε E2α’ helix only impacts activation by RhoA. PLCε is activated by RhoA^G14V^, a constitutively active mutant of Rap1A (Rap1A^Q63E)^, and the Gβγ heterodimer. PLCε Δ11275-1289 is also responsive to the G proteins. However, PLCε and PLCε Δ11275-1289 only differ in their fold-activation over basal upon co-transfection with RhoA^G14V^. (*Top left*) At least three independent experiments from independent transfections were performed for each variant, and data is shown as the average of triplicate measurements ± SD. Data was analyzed using an unpaired, one-tailed t-test with Welch’s correction comparing the basal and G protein-stimulated activities of each variant. For PLCε and RhoA^G14V^, ****p<0.0001, with Rap1A^Q63E^ *p<0.0188, and with Gβγ *p<0.0241. For PLCε Δ11275-1289 and RhoA^G14V^ **p<0.0017, with Rap1A^Q63E^ *p<0.0140, and with Gβγ **p<0.0032. (*Top right*) Fold activation was calculated by dividing the maximum activity measured with a G protein by the basal activity of either PLCε or PLCε Δ11275-1289. The only significant difference in the ability of the G proteins to activate either PLCε or PLCε Δ11275-1289 was observed with RhoA^G14V^. Data was analyzed using an unpaired, one-tailed t-test with Welch’s correction comparing the fold activations of PLCε and PLCε Δ11275-1289 for each G protein. **p<0.0073. Representative western blots are shown below, with empty pCMV vector (EV) and β-actin used as loading controls. PLCε variants express a C-terminal FLAG tag and are detected using an anti-FLAG antibody, RhoA^G14V^ and Rap1A^Q63E^ contain N-terminal HA tags, and were detected using an anti-HA antibody. Endogenous and Avi-tagged Gβγ were detected using an antibody against the Gβ subunit.

**Supplemental Figure 12.**
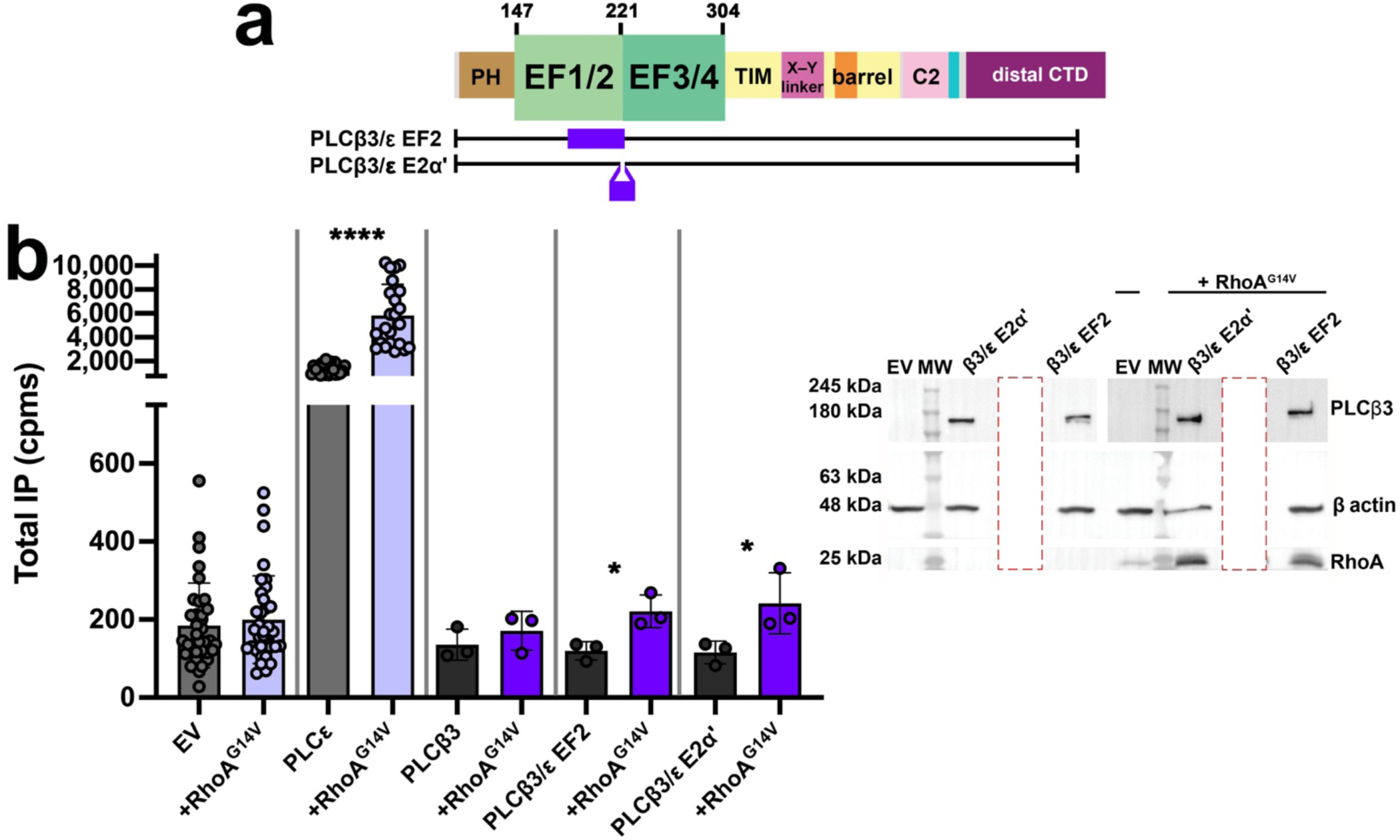
The PLCε E2α’ helix confers sensitivity to RhoA. **(a)** Schematic of the PLCβ3/ε chimeras. In PLCβ3/ε EF2, E2α-E3α (residues 183-221) are replaced with PLCε F2α-E2α’ (residues 1196-1284). In PLCβ3/ε E2α’, the PLCε E2α’ helix and linker to EF3 are inserted between the PLCβ3 EF1/2-EF3/4 subdomains (residues 221-222). **(b)** At least three independent experiments from independent transfections were carried out for each variant, and data is shown as the average of triplicate measurements ± SD. Data was analyzed using an unpaired, one-tailed t-test with Welch’s correction comparing the basal and RhoA-stimulated activities of each variant. ****p<0.0001; *p=0.0167 for PLCβ3/ε EF2, *p=0.0472 for PLCβ3/ε E2α’. Representative western blot of the PLCβ3/ε chimeras are shown, with empty pCMV vector (EV) and β-actin used as loading controls. PLCβ3 chimeras, β-actin, and HA-RhoA were incubated with rabbit anti-PLCβ3, mouse anti-β-actin, and mouse anti-HA primary antibodies, respectively.

## Notes

### Competing Interest Statement

The authors have declared no competing interest.

### Summary of Updates

cryo-EM data collection, processing, and structure validation, new supplemental figures, edits for clarity in the main text and supplement.

## REFERENCES

1. Kadamur, G. & Ross, E.M. Mammalian phospholipase C. Annu Rev Physiol 75, 127–54 (2013).

2. Smrcka, A.V., Brown, J.H. & Holz, G.G. Role of phospholipase Cε in physiological phosphoinositide signaling networks. Cell Signal 24, 1333–43 (2012).

3. Muralidharan, K., Van Camp, M.M. & Lyon, A.M. Structure and regulation of phospholipase Cbeta and epsilon at the membrane. Chem Phys Lipids 235, 105050 (2021).

4. Smrcka, A.V. Regulation of phosphatidylinositol-specific phospholipase C at the nuclear envelope in cardiac myocytes. J Cardiovasc Pharmacol 65, 203–10 (2015).

5. Oestreich, E.A. et al. Epac and phospholipase Cε regulate Ca^2+^ release in the heart by activation of protein kinase Cε and calcium-calmodulin kinase II. J Biol Chem 284, 1514–22 (2009).

6. Oestreich, E.A. et al. Epac-mediated activation of phospholipase Cε plays a critical role in beta-adrenergic receptor-dependent enhancement of Ca^2+^ mobilization in cardiac myocytes. J Biol Chem 282, 5488–95 (2007).

7. Wang, H. et al. Phospholipase C ε modulates β-adrenergic receptor-dependent cardiac contraction and inhibits cardiac hypertrophy. Circ Res 97, 1305–13 (2005).

8. Zhang, L., Malik, S., Kelley, G.G., Kapiloff, M.S. & Smrcka, A.V. Phospholipase C ε scaffolds to muscle-specific A kinase anchoring protein (mAKAPβ) and integrates multiple hypertrophic stimuli in cardiac myocytes. J Biol Chem 286, 23012–21 (2011).

9. Zhang, L. et al. Phospholipase cepsilon hydrolyzes perinuclear phosphatidylinositol 4-phosphate to regulate cardiac hypertrophy. Cell 153, 216–27 (2013).

10. Nash, C.A., Brown, L.M., Malik, S., Cheng, X. & Smrcka, A.V. Compartmentalized cyclic nucleotides have opposing effects on regulation of hypertrophic phospholipase Cepsilon signaling in cardiac myocytes. J Mol Cell Cardiol (2018).

11. Wei, W. & Smrcka, A.V. Internalized β2-Adrenergic Receptors Oppose PLC-Dependent Hypertrophic Signaling. Circulation Research 135(2024).

12. Malik, S. et al. G protein betagamma subunits regulate cardiomyocyte hypertrophy through a perinuclear Golgi phosphatidylinositol 4-phosphate hydrolysis pathway. Mol Biol Cell 26, 1188–98 (2015).

13. Means, C.K. & Brown, J.H. Sphingosine-1-phosphate receptor signalling in the heart. Cardiovasc Res 82, 193–200 (2009).

14. Xiang, S.Y., et al. PLCepsilon, PKD1, and SSH1L transduce RhoA signaling to protect mitochondria from oxidative stress in the heart. Sci Signal 6, ra108 (2013).

15. Yung, B.S. et al. Selective coupling of the S1P 3 receptor subtype to S1P-mediated RhoA activation and cardioprotection. Journal of Molecular and Cellular Cardiology 103, 1–10 (2017).

16. Xiang, S.Y. et al. RhoA protects the mouse heart against ischemia/reperfusion injury. Journal of Clinical Investigation 121, 3269–3276 (2011).

17. Brand, C.S., Tan, V.P., Brown, J.H. & Miyamoto, S. RhoA regulates Drp1 mediated mitochondrial fission through ROCK to protect cardiomyocytes. Cellular Signalling 50, 48–57 (2018).

18. Samassekou, K., et al. (2024).

19. Bunney, T.D. et al. Structural and mechanistic insights into ras association domains of phospholipase C ε. Mol Cell 21, 495–507 (2006).

20. Sieng, M. et al. Functional and structural characterization of allosteric activation of phospholipase Cepsilon by Rap1A. J Biol Chem 295, 16562–16571 (2020).

21. Hicks, S.N. et al. General and versatile autoinhibition of PLC isozymes. Mol Cell 31, 383–94 (2008).

22. Lyon, A.M., Begley, J.A., Manett, T.D. & Tesmer, J.J. Molecular Mechanisms of Phospholipase C beta3 Autoinhibition. Structure 22, 1844–1854 (2014).

23. Seifert, J.P. et al. RhoA activates purified phospholipase C-ε by a guanine nucleotide-dependent mechanism. J Biol Chem 279, 47992–7 (2004).

24. Kelley, G.G., Kaproth-Joslin, K.A., Reks, S.E., Smrcka, A.V. & Wojcikiewicz, R.J. G-protein-coupled receptor agonists activate endogenous phospholipase Cε and phospholipase Cβ3 in a temporally distinct manner. J Biol Chem 281, 2639–48 (2006).

25. Kelley, G.G., Reks, S.E. & Smrcka, A.V. Hormonal regulation of phospholipase Cε through distinct and overlapping pathways involving G_12_ and Ras family G-proteins. Biochem J 378, 129–39 (2004).

26. Seifert, J.P., Zhou, Y., Hicks, S.N., Sondek, J. & Harden, T.K. Dual activation of phospholipase C-ε by Rho and Ras GTPases. J Biol Chem 283, 29690–8 (2008).

27. Dusaban, S.S. et al. Phospholipase C{varepsilon} links G protein-coupled receptor activation to inflammatory astrocytic responses. Proc Natl Acad Sci U S A 110, 3609–14 (2013).

28. Wing, M.R., Snyder, J.T., Sondek, J. & Harden, T.K. Direct activation of phospholipase C-ε by Rho. J Biol Chem 278, 41253–8 (2003).

29. de Rubio, R.G. et al. Phosphatidylinositol 4-phosphate is a major source of GPCR-stimulated phosphoinositide production. Science Signaling 11(2018).

30. Rugema, N.Y. et al. Structure of phospholipase Cepsilon reveals an integrated RA1 domain and previously unidentified regulatory elements. Commun Biol 3, 445 (2020).

31. Citro, S. et al. Phospholipase Cε is a nexus for Rho and Rap-mediated G protein-coupled receptor-induced astrocyte proliferation. Proc Natl Acad Sci U S A 104, 15543–8 (2007).

32. Lyon, A.M., Dutta, S., Boguth, C.A., Skiniotis, G. & Tesmer, J.J. Full-length Galpha(q)-phospholipase C-beta3 structure reveals interfaces of the C-terminal coiled-coil domain. Nat Struct Mol Biol 20, 355–62 (2013).

33. Esquina, C.M. et al. Intramolecular electrostatic interactions contribute to phospholipase Cbeta3 autoinhibition. Cell Signal, 109349 (2019).

34. Hudson, B., Jessup, R.E., Prahalad, K.K. & Lyon, A.M. Galphaq and the Phospholipase Cbeta3 X-Y Linker Regulate Adsorption and Activity on Compressed Lipid Monolayers. Biochemistry (2019).

35. Garland-Kuntz, E.E. et al. Direct observation of conformational dynamics of the PH domain in phospholipases C and beta may contribute to subfamily-specific roles in regulation. J Biol Chem 293, 17477–17490 (2018).

36. Dvorsky, R., Blumenstein, L., Vetter, I.R. & Ahmadian, M.R. Structural Insights into the Interaction of ROCKI with the Switch Regions of RhoA. Journal of Biological Chemistry 279, 7098–7104 (2004).

37. Chen, Z. et al. Activated RhoA Binds to the Pleckstrin Homology (PH) Domain of PDZ-RhoGEF, a Potential Site for Autoregulation. Journal of Biological Chemistry 285, 21070–21081 (2010).

38. Kristelly, R., Gao, G. & Tesmer, J.J.G. Structural Determinants of RhoA Binding and Nucleotide Exchange in Leukemia-associated Rho Guanine-Nucleotide Exchange Factor. Journal of Biological Chemistry 279, 47352–47362 (2004).

39. Essen, L.O., Perisic, O., Cheung, R., Katan, M. & Williams, R.L. Crystal structure of a mammalian phosphoinositide-specific phospholipase C δ. Nature 380, 595–602 (1996).

40. Chen, J.E., Huang, C.C. & Ferrin, T.E. RRDistMaps: a UCSF Chimera tool for viewing and comparing protein distance maps. Bioinformatics 31, 1484–1486 (2015).

41. Madukwe, J.C., Garland-Kuntz, E.E., Lyon, A.M. & Smrcka, A.V. G protein betagamma subunits directly interact with and activate phospholipase Cepsilon. J Biol Chem (2018).

42. Chen, C.-L. et al. Molecular basis for Gβγ-mediated activation of phosphoinositide 3-kinase γ. Nature Structural & Molecular Biology 31, 1198–1207 (2024).

43. Falzone, Maria E. & MacKinnon, R. Gβγ activates PIP2 hydrolysis by recruiting and orienting PLCβ on the membrane surface. Proceedings of the National Academy of Sciences 120(2023).

44. Thompson, R.F., Iadanza, M.G., Hesketh, E.L., Rawson, S. & Ranson, N.A. Collection, pre-processing and on-the-fly analysis of data for high-resolution, single-particle cryo-electron microscopy. Nature Protocols 14, 100–118 (2018).

45. Liebschner, D. et al. Macromolecular structure determination using X-rays, neutrons and electrons: recent developments in Phenix. Acta Crystallographica Section D Structural Biology 75, 861–877 (2019).

46. Adams, P.D. et al. PHENIX: a comprehensive Python-based system for macromolecular structure solution. Acta Crystallogr D Biol Crystallogr 66, 213–21 (2010).

47. Casañal, A., Lohkamp, B. & Emsley, P. Current developments in Coot for macromolecular model building of Electron Cryo-microscopy and Crystallographic Data. Protein Science 29, 1055–1064 (2020).

48. Terashi, G., Wang, X., Maddhuri Venkata Subramaniya, S.R., Tesmer, J.J.G. & Kihara, D. Residue-wise local quality estimation for protein models from cryo-EM maps. Nature Methods 19, 1116–1125 (2022).

49. Prisant, M.G., Williams, C.J., Chen, V.B., Richardson, J.S. & Richardson, D.C. New tools in MolProbity validation: CaBLAM for CryoEM backbone, UnDowser to rethink “waters,” and NGL Viewer to recapture online 3D graphics. Protein Science 29, 315–329 (2019).

